# Gut microbes mediate the synergistic effects of dietary cholesterol and saturated fat in driving fibrosing MASH

**DOI:** 10.1101/2025.07.16.665145

**Authors:** Jake B. Hermanson, Samar A. Tolba, Md Amran Gazi, Evan A. Chrisler, Manpreet Kaur, Ashley M. Sidebottom, Yongjun Liu, Guillermo Martinez-Boggio, Lauren N. Lucas, Daniel Amador-Noguez, Federico E. Rey, Vanessa A. Leone

## Abstract

Metabolic dysfunction-associated steatotic liver disease (MASLD) affects approximately one-third of the global population and can progress to metabolic dysfunction-associated steatohepatitis (MASH) with fibrosis, increasing risk of cirrhosis, hepatocellular carcinoma, and mortality. Gut microbes driven by diets high in saturated fat, simple sugar, and cholesterol contribute to disease progression, yet underlying mechanisms remain undefined. We explored the independent and synergistic effects of dietary saturated fat and cholesterol on MASH development using specific pathogen-free (SPF) and germ-free (GF) mice. We demonstrate that 1) both dietary cholesterol and saturated fat are required to induce fibrosing MASH in SPF mice, whereas GF mice are protected, 2) saturated fat and cholesterol individually alter gut microbial membership, potentially via altered bile acid metabolism, while their combination promotes a distinct composition, including an increase in *Parasutterella* spp. which correlates with hepatic fibrosis, and 3) diluted cecal contents from SPF, but not GF, mice fed high-fat, high-cholesterol diets are enriched in deoxycholic acid and activate human hepatic stellate cells *in vitro*, suggesting a mechanistic link between dietary lipid-induced microbiota and liver fibrogenesis. These findings reveal how specific Western dietary components shape the gut microbiota and contribute to hepatic liver fibrosis via stellate activation, offering potential targets for therapeutic intervention in MASLD/MASH.

## INTRODUCTION

Metabolic dysfunction-associated steatotic liver disease (MASLD, formerly nonalcoholic fatty liver disease [NAFLD]^1^) is the most common chronic liver disease, impacting nearly a third of the population worldwide^2^. Approximately 15-20% of MASLD patients progress to Metabolic dysfunction-associated steatohepatitis (MASH), which can lead to cirrhosis and hepatocellular carcinoma (HCC)^3,4^. MASH will soon become the leading indication for liver transplantation^5^. Hepatic fibrosis, a hallmark of advanced MASH, associates with worse disease outcomes. Higher fibrosis stages are associated with ∼10 times greater risk of liver-related mortality^6^. Despite growing disease burden, treatment options remain limited. Resmetirom (marketed as Rezdiffra™), which was recently approved by the U.S. Food and Drug Administration as a first-in-class therapy for fibrosing MASH, achieved primary endpoints for fibrosis stage improvement in only 25-29% of patients, and its long-term efficacy is not fully understood^7^. These limitations underscore the need for additional therapeutic interventions.

MASLD and MASH are heterogeneous diseases, shaped by complex and dynamic interactions among genetic and environmental factors, including diet and the trillions of gut microbes that reside within the intestine^8^. Deciphering the mechanistic connections between these factors is essential to advance our understanding of MASLD and MASH disease etiology for the development of effective, targeted interventions. Dietary cholesterol has emerged as a potent disease driver of MASLD and MASH pathogenesis. In both humans^9–15^ and preclinical animal models^16–18^, elevated dietary cholesterol strongly associates with disease prevalence and severity. For instance, in a large cohort of ∼215,000 individuals, consuming ≥121.4 mg cholesterol per day increased MASLD risk, particularly among those with cirrhosis^10^. In mice, dietary cholesterol is required to induce fibrosing MASH; while a high-fat diet (HFD) alone induces simple steatosis^17,18^, incorporating 0.2% cholesterol (wt/wt) drives inflammation, fibrosis, and progression to HCC^19^. It has been posited that cholesterol exacerbates steatosis and lipotoxicity, which in turn promote hepatic inflammation and eventually fibrosis^20^. Despite this, cholesterol’s role in disrupting the gut-liver axis, particularly through its interaction with gut microbes, remains poorly understood. Further, disentangling the specific contributions of dietary cholesterol and saturated fat to these disruptions is also challenging in humans as both nutrients co-occur in animal products such as red meat and eggs.

Gut microbes are known to influence metabolic diseases, including MASLD/MASH. Germ-free (GF) mice are generally protected from diet-induced obesity and liver pathology^21,22^, whereas microbial transplantation from affected Specific pathogen-free (SPF) mice can transfer pathological features to GF recipients^19,23^. For example, long-term feeding of high saturated fat and cholesterol diet in mice leads to sequential development of steatosis, fibrosis, and HCC relative to chow or HFD^19^. This diet altered serum metabolites, increasing taurocholic acid (TCA) and decreasing 3-indolepropionic acid (IPA). These changes were accompanied by shifts in gut microbes, including enrichment of *Mucispirillum* and *Desulfovibrio* and depletion of *Bifidobacteria*. *In vitro*, TCA promoted hepatic lipid accumulation, while IPA led to reductions, suggesting gut microbe-derived metabolites mediate key pathways in MASLD/MASH progression^19^. While these findings implicate microbial metabolites in steatosis, their contribution to fibrogenesis, as well as the distinct roles of dietary cholesterol vs. saturated fat, remains unclear. Gut microbes can enzymatically transform dietary cholesterol into non-absorbable coprostanol and other metabolites^24–27^, thereby altering homeostasis of host cholesterol metabolism. Studies by Le et al. and Yao et al. revealed certain bacteria, e.g., *Bifidobacterium pseudolongum*, *Enterococcus*, and *Parabacteroides*, can contribute to either direct metabolism of exogenous cholesterol via various enzymes, e.g., sulfotransferases, or perform uptake and assimilation^26,27^. These studies and others underscore the complex interplay between dietary lipids and gut microbes in liver disease pathogenesis. In this study, we sought to determine how dietary cholesterol vs. saturated fat independently and synergistically reshape gut microbiota composition in the context of MASLD/MASH progression using a murine model. We hypothesized that while each of these components alters gut microbiota and host physiology in unique ways, their combination is required to induce fibrosing MASH in SPF mice, whereas their GF counterparts will remain largely protected against disease. We show that the combination of dietary cholesterol and saturated fat is essential to drive gut microbiota imbalances and disrupted bile acid metabolism that contribute to hepatic stellate cell (HSC) activation and advancement to fibrosing MASH.

## MATERIALS AND METHODS

### Animals

8-week-old male Specific pathogen-free (SPF) C57Bl/6J mice were purchased from the Jackson Laboratories (barrier facility MP15) and maintained on a 12:12 hour light:dark cycle. From 8 to 16 weeks of age, mice were housed 4/cage and provided autoclaved aspen shavings (Waldschmidt & Sons, Madison, WI), water and LabDiet® 5k67 chow. Bedding was mixed twice weekly across all cages to normalize microbiome composition as previously described^28^. Male germ-free (GF) C57Bl/6 mice were bred in flexible film gnotobiotic isolators (CBC Clean, Inc., Madison, WI) at the University of Wisconsin-Madison Gnotobiotic facility and provided autoclaved aspen shavings, water, and LabDiet® 5K67 chow from 8 to 16 weeks of age. GF status was confirmed via 16S rRNA PCR on freshly collected fecal pellets weekly and via routine fecal cultivation under anaerobic and aerobic conditions. At 16 weeks of age, both SPF and GF mice were housed 2/cage and randomly assigned to one of six diets: low-fat (LF), low-fat+high-cholesterol (LFHC), low-fat+very high-cholesterol (LFVHC), high-fat (HF), high-fat+high cholesterol (HFHC), or high-fat+very high-cholesterol (HFVHC) (**Table S1**). Fresh feces and plasma (via submandibular vein) were collected at baseline and every 4 weeks from SPF mice and every 8 weeks from GF mice and stored at −80°C. Body weight as well as food and water consumption were measured weekly. After 8 and 24 weeks on diet, mice were euthanized via CO_2_ asphyxiation. Portal and cardiac blood were collected into heparinized tubes, and 5 x 5 x 5 mm liver sections were prepared for histology. The remaining liver tissue was flash-frozen and stored at −80°C. Epididymal, mesenteric, retroperitoneal, and inguinal adipose tissue as well as cecal luminal contents were weighed, flash-frozen, and stored at −80°C.

### ALT Measurement

∼200µL blood was collected via submandibular vein into heparin-coated microfuge tubes followed by centrifugation (10,000xG) for 10 minutes at 4°C to obtain plasma. Plasma was diluted ¼ in phosphate buffered saline (PBS) and Alanine transaminase (ALT) activity was measured colorimetrically on a Catachem Well-T AutoAnalyzer using the ALT Dual Kit (Catachem, Oxford, CT).

### Total Bile Acid Quantification

Fecal pellets were weighed, dried overnight at 55°C in glass tubes, and reweighed to obtain dry weight. 2mL Folch solution (2:1 chloroform:methanol) was added, and samples were incubated in a 60°C water bath while shaking (∼40rpm) for 30 minutes. Samples were centrifuged at 1500xG for 10 minutes. The lower chloroform phase was transferred to a new 1.5mL centrifuge tube and evaporated under a N_2_ gas stream. The resulting lipids were resuspended in 100µL 1% Triton X-100 in EtOH. Total bile acid concentration in fecal lipid extracts and cell-free cecal homogenates were determined using the Crystal Chem Mouse Total Bile Acids Assay Kit (Crystal Chem, Elk Grove Village, IL) according to manufacturer’s instructions.

### Endotoxin (LPS) Quantification

LPS concentrations in cell-free cecal homogenates were measured using the Pierce Chromogenic Endotoxin Quant Kit (Thermo Fisher Scientific, Waltham, MA) according to manufacturer’s instructions.

### Liver Histology

For Oil-Red-O staining, liver sections were cryopreserved in O.C.T. (Tissue-Tek) and 10µm sections were cut using a Leica cryotome. Sections were fixed with 4% paraformaldehyde (PFA) for 15 minutes and stained with Oil-Red-O (lipid droplets) and hematoxylin (nuclei)^29^. Liver histology of formalin-fixed tissue was performed at the UW-Madison Translational Research Initiatives in Pathology (TRIP) lab. Briefly, following fixation in 4% neutral buffered formalin for 24 hours, tissue was transferred to 70% EtOH, paraffin-embedded, and 5µm sections were cut via a microtome. Following deparaffinization, sections were stained with H&E, Picrosirius Red, or Masson’s Trichrome. NAFLD Activity Score (NAS) was determined on H&E sections by a trained pathologist, blinded to subject treatment, according to Kleiner et al^30^ based on hepatic steatosis, lobular inflammation, and hepatocyte ballooning. Lipid droplets (Oil-Red-O) and collagen deposition (Picrosirius Red, Masson’s Trichrome) were quantified using ImageJ 2 (v 1.53a). Briefly, for Oil-Red-O and Picrosirius Red stained sections, raw images were split into red, green, and blue greyscale channels. The green channel was selected, and a color threshold was set to highlight stained areas. For Masson’s Trichrome, raw images were split into a “Lab” greyscale stack, the “b” channel was selected, and a color threshold was set to highlight stained areas.

### RNA Extraction and Quantitative Real-Time Reverse Transcription PCR (qRT-PCR)

RNA was extracted from ∼15 mg of liver tissue using TRIzol reagent and chloroform as previously described^31^. cDNA was prepared using the iScript™ gDNA Clear cDNA Synthesis Kit (Bio-Rad, Hercules, CA) following the manufacturer’s protocol. cDNA was combined with SYBR Green qPCR Master Mix (Bio-Rad, Hercules, CA) and forward and reverse primers (**Table S2**) and quantification of each gene was obtained on a CFX384 Real-Time PCR Detection System (Bio-Rad, Hercules, CA). Data were normalized using *Glyceraldehyde-3-phosphate dehydrogenase* (*Gapdh*) as the housekeeping gene and presented as 2^(-ΔΔCt)^, with Week 8 SPF LF set as the control.

### 16S rRNA Gene Amplicon Sequencing and Analysis

DNA was extracted from feces and cecal contents as previously described^28^. The V4 region of the 16S rRNA gene was amplified using 515F-806R primers (**Table S1**). PCR amplification was performed at 94°C for 3 minutes followed by 40 cycles at 94°C (45 seconds), 50°C (60 seconds), and 72°C (90 seconds). Paired-end reads (150 x 150bp) of the resulting amplicons were sequenced on an Illumina MiSeq at Argonne National Laboratory. A total of 13,949,738 (fecal samples) and 2,804,624 (cecal samples) raw reads were obtained, with an average value of 35,952 (fecal samples) or 36,902 (cecal samples) reads per sample. Paired-end demultiplexed reads were imported and filtered utilizing Quantitative Insights Into Microbial Ecology (QIIME2, 2024.2)^32^ and trimmed to 120bp. Divisive amplicon denoising algorithm (DADA2)^33^ was used to filter and denoise the imported demultiplexed sequences (via q2-dada2), where a total of 12,619,444 (fecal samples) and 2,453,434 (cecal samples) reads passed quality checks with an average of 32,524 (fecal samples) and 32,282 (cecal samples) reads/sample. All samples were rarified to a sequencing depth of 15,000 sequences per sample. α- and β-diversity metrics and Principal Coordinate Analysis (PCoA) were performed using the q2-diversity plugin in R. Taxonomy was assigned to amplicon sequence variants (ASVs) using the Silva-138 99% reference sequences via the q2-feature-classifier.

### Cell Culture

LX-2 human immortalized hepatic stellate cells (Sigma-Aldrich, St. Louis, MO) were grown in Dulbecco’s Modified Eagle Medium (DMEM, Thermo Fisher Scientific, Waltham, MA) supplemented with 2% fetal bovine serum (FBS, Thermo Fisher Scientific, Waltham, MA), 1000 U/mL penicillin, 1000 µg/mL streptomycin (Pen/Strep, Thermo Fisher Scientific, Waltham, MA), and 20 mM glutamine (Thermo Fisher Scientific, Waltham, MA) in a cell culture incubator set to 37°C, 5% CO_2_, 90-95% relative humidity^34,35^. Prior to experiments, cells were seeded in 12-well (∼1×10^5^ cells/well) or 6-well (∼3×10^5^ cells/well) tissue culture-treated cell culture plates (Corning, Corning, NY) and incubated for 24 hours or until ∼80% confluent. Serum starvation was then performed to normalize the cell cycle across all wells by replacing media with 0.2% FBS media for 24 hours.

Cell-free cecal homogenates were prepared by suspending cecal contents in sterile PBS at 50 mg/mL. Samples were homogenized using a pellet pestle cordless motor (Thermo Fisher Scientific, Waltham, MA) for three rounds of 30 seconds. Homogenates were centrifuged at 10,000xG (4°C), and the supernatant was passed through a 0.22 µm filter and stored at −80°C. Once LX-2 cells reached ∼80% confluence, wells were washed twice with sterile PBS and pre-warmed media containing cell-free cecal homogenate (10% vol/vol) were added. In separate experiments, deoxycholic acid (Thermo Fisher Scientific, Waltham, MA was added to growth media at the indicated concentrations. After four hours, media was removed, cells were washed twice with sterile PBS, and RNA was extracted using a RNeasy Mini Kit (Qiagen, Germantown, MD) according to manufacturer’s instructions. cDNA preparation and qRT-PCR were performed as described above.

### Fecal Bile Acid Measurements

Lipidomics were performed at the Duchossois Family Institute at the University of Chicago. Metabolites were extracted from fecal samples by adding 1mL of 80% methanol (spiked with internal standards) per 100mg feces and homogenized at 4°C on a Bead Mill 24 Homogenizer (1.6m/s, six 30-second cycles, 5 seconds off between samples). Homogenates were centrifuged at 20,000xG (−10°C, 15 minutes) and supernatant was collected. 75µL of supernatant was added to autosampler vials, dried via nitrogen stream (30 L/min on top; 1 L/min on bottom, 30°C), and resuspended with a thermomixer (4°C, 1000rpm, 15 minutes) in 750µL of 50% methanol. Insoluble debris was removed via centrifugation (4°C, 20000xG, 15 minutes) and the supernatant was transferred to an autosampler vial.

Metabolites were analyzed using a liquid chromatograph (Agilent 1290 Infinity II) coupled to a quadrupole time-of-flight (QTOF) mass spectrometer (Agilent 6546) in negative mode with an Agilent Jet Stream Electrospray Ionization source. 5µL of sample was injected into an XBridge© BEH C18 Column (3.5 µm, 2.1 x 100mm; Waters Corporation) with an XBridge© BEH C18 guard (Waters Corporation) at 45°C. Elution began with 72% A (Water, 0.1% formic acid) and 28% B (Acetone, 0.1% formic acid) at 0.4 mL/minute for 1 minute and increased to 33% B over a 5-minute period, then increased to 65% B over a 14-minute period. Flow rate was increased to 0.6 mL/minute and increased to 98% B over a 30-second period. This condition was held constant for 3.5 minutes. Flow rate was decreased to 0.4 mL/minute and 28% B for 3 minutes to re-equilibrate. Electrospray ionization conditions were set (capillary voltage: 3.5 kV, nozzle voltage: 2kV, detection window: 100-1700 *m/z*) with continuous infusion of reference masses (Agilent ESI TOF Biopolymer Analysis Reference Mix). A calibration curve (10-point) was used for quantitation and data was analyzed using MassHunter Profinder Analysis (v B.10, Agilent Technologies) and validated by comparison to authentic standards. For bile acids that were outside the range of the standard curve, normalized peak areas were calculated by dividing the raw peak areas of the target analytes by the averaged raw peak areas of the internal standards. Z-scores were determined for each sample within a given bile acid by the following formula:

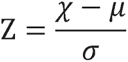

Where *χ* is the normalized peak area of the sample, *μ* is the mean peak area for a given bile acid across all samples and *σ* is the standard deviation for a given bile acid across all samples.

### Cecal Homogenate Bile Acid Measurements

Cecal homogenates were analyzed using an ultra-high pressure liquid chromatography-tandem mass spectrometry (uHPLC-MS/MS) system consisting of a ThermoScientific Vanquish uHPLC system coupled to a heated electrospray ionization (HESI; using negative polarity) and hybrid quadrupole high resolution mass spectrometer (Q Exactive Orbitrap; Thermo Scientific). Settings for the ion source were: auxiliary gas flow rate of 10, sheath gas flow rate of 30, sweep gas flow rate of 1, 2.5 kV spray voltage, 320°C capillary temperature, 300°C heater temperature, and S-lens RF level of 50. Nitrogen was used as nebulizing gas by the ion trap source. Liquid chromatography (LC) separation was achieved using a Waters Acquity UPLC BEH C18 column with 1.7 μm particle size, 2.1 x 100 mm in length. Solvent A was water with 10 mM ammonium acetate adjusted to pH 6.0 with acetic acid. Solvent B was 100% methanol. The total run time was 31.5 min with the following gradient: a 0 to 24 min gradient from 30% solvent B (initial condition) to 100% solvent B; held 5 min at 100% solvent B; dropped to 30% solvent B for 2.5 min re-equilibration to initial condition. The flow rate was 200 μL/min throughout. Other LC parameters were as follows: autosampler temperature, 4°C; injection volume, 10 μL; column temperature 50°C. The MS method performed a full MS1 full-scan (290 to 1000 m/z) together with a series of PRM (parallel reaction monitoring) scans. Untargeted experimental MS data were converted to the mzXML format and used for targeted bile acid identification using El-MAVEN and matching sample peaks to standard peaks^36^. Bile acids were quantified using 8-point external standard curves, with each bile acid ranging from .03125 to 4 μM, allowing for conversion of raw signal to μM concentration. Samples were run at both 1/10 and 1/100 dilutions for all bile acid measurements to fall within the external standard signal range. The detection limit was below 0.01 μM for all bile acids. The threshold for reported core bile acid transformations was 0.01 μM. Standards were purchased from Avanti Polar Lipids and dissolved and stored in methanol at −80 °C. See **Table S3** for bile acid standard names and structural features.

### Statistical Analysis

Statistical analyses were performed using GraphPad Prism (v 10.2.3) or R (v 4.4.2). Unless otherwise stated, data represent means ± standard error of the mean (SEM). Total caloric intake, food intake, water intake, and body weight gain over time (**Figure 1B; S1A**) were analyzed using a 4-way repeated measures (RM) ANOVA following a 2 x 2 x 2 x K factorial design (Factors: Cholesterol, Fat, Microbes, Time) where K = weeks on diet and accounts for repeated measures within the same mouse and accounts for subject effect. If significant interaction or main effects were observed (*P*<0.05), 3-way RM ANOVA was performed within SPF and GF groups (2 x 2 x K factorial; Factors: Cholesterol, Fat, Time).

**Figure 1.**
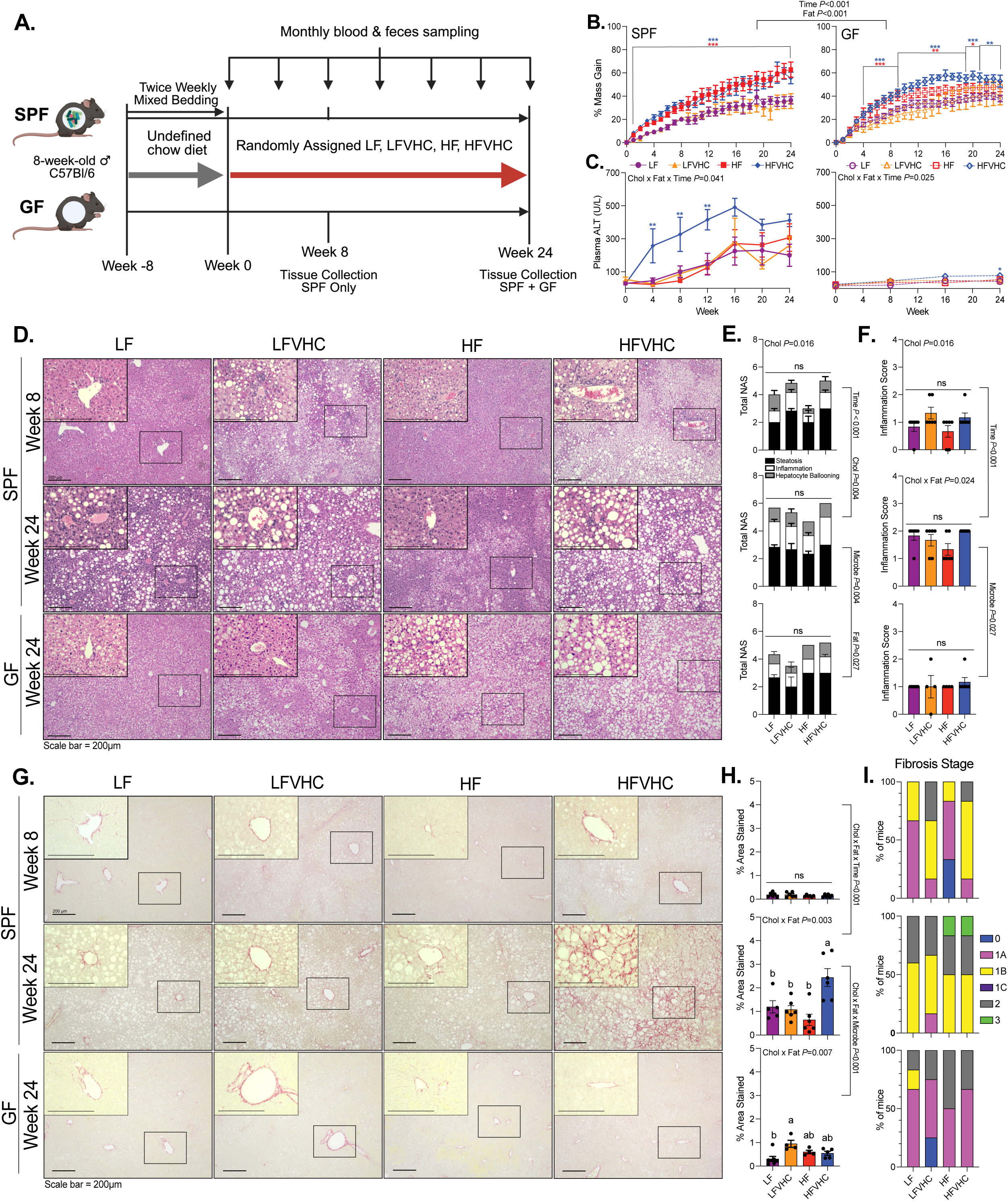
HFVHC diet drives fibrosing MASH in SPF but not GF mice. **A)** Experimental schematic. Created on Biorender.com. **B)** Percent body mass gain over time relative to baseline in SPF (left) and GF (right) mice. Data analyzed via 4-way repeated measures (RM) ANOVA (factors: Cholesterol, Fat, Microbes, Time) followed by 3-way ANOVA (factors: Cholesterol, Fat, Time) within SPF and GF groups and Tukey’s multiple comparisons within timepoint. **C)** Plasma Alanine transaminase (ALT) levels, analyzed via 3-way RM ANOVA (factors: Cholesterol, Fat, Time) within SPF and GF groups followed by Tukey’s multiple comparisons within timepoint. For panels B and C, \**P*<0.05, \*\**P*<0.01, \*\*\**P*<0.005. Asterisk color represents which group is significantly different from LF. **D)** Representative H&E-stained liver sections; scale bar=200µm. Inset images (400x) correspond to boxed regions in 100x images. **E)** Total NAFLD activity score (NAS) split by components and **F)** lobular inflammation scores based on H&E sections. **G)** Representative picrosirius red-stained liver sections; scale bar=200µm. Inset images (400x) correspond to boxed regions in 100x images. **H)** Quantification of percent area stained (indicating collagen). Data analyzed via 3-way ANOVA, comparing SPF Week 8 vs. SPF Week 24 (factors: Cholesterol, Fat, Time) and SPF Week 24 vs. GF Week 24 (factors: Cholesterol, Fat, Microbes), followed by Tukey’s multiple comparisons. Bars with the same letter are not significantly different (*P*>0.05). **I)** Fibrosis stage based on picrosirius red staining. All data represent means ± SEM, unless noted otherwise.

ALT data (**Figure 1C**) were analyzed using a 3-way RM ANOVA following a 2 x 2 x K factorial design (Factors: Cholesterol, Fat, Time) within SPF and GF groups. Fecal α-diversity indices (**Figure S2A**) and relative abundance of fecal ASVs (**Figure 3C,D**) within SPF mice were analyzed via 3-way RM ANOVA following a 2 x 2 x K factorial design (Factors: Cholesterol, Fat, Time). If significant interactive or main effects were observed (*P*<0.05), 2-way ANOVA based on a 2 x 2 factorial design (Factors: Cholesterol, Fat) was performed within timepoint.

Histological analyses (**Figure 1E,F,H; Figure S1E,G**) as well as adipose and liver mass (**Figure S1B,C**) were analyzed via one of two models:

Model 1 in SPF mice = 3-way ANOVA (2 x 2 x 2 factorial; Factors: Cholesterol, Fat, Time), where “Time” accounted for samples collected at 8 and 24 weeks. If significant interactive or main effects were observed (*P*<0.05), 2-way ANOVA based on a 2 x 2 factorial design (Factors: Cholesterol, Fat) was performed within timepoint.

Model 2 in SPF and GF mice = 3-way ANOVA (2 x 2 x 2 factorial; Factors: Cholesterol, Fat, Microbes) where “Microbe” accounted for samples collected at 24 weeks from SPF and GF mice. If significant interactive or main effects were observed (*P*<0.05), 2-way ANOVA based on a 2 x 2 factorial design (Factors: Cholesterol, Fat) was performed within SPF and GF groups.

Murine liver gene expression (**Figure 2B,C**), fecal BA (**Figure 4A,B,G-L**), and cecal α-diversity (**Figure S2B**), were analyzed via 3-way ANOVA following a 2 x 2 x 2 factorial design (Factors: Cholesterol, Fat, Time) in SPF mice only. If a significant interaction or main effect was detected (*P*<0.05), posthoc Tukey’s multiple comparisons were performed.

**Figure 2.**
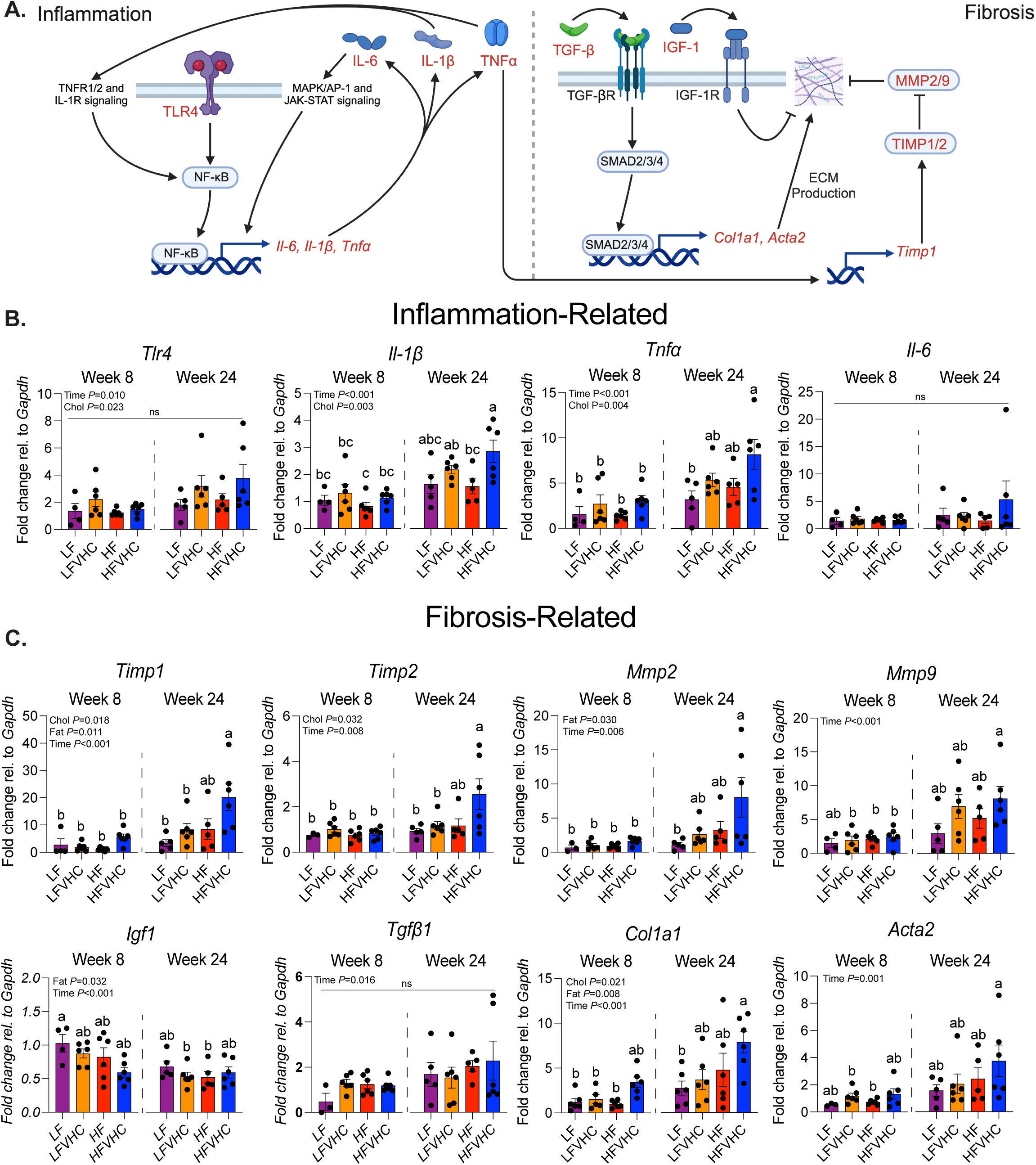
Dietary cholesterol enhances expression of inflammation- and fibrosis-related genes in SPF mice. **A)** Schematic of key signaling pathways involved in inflammation and fibrosis in MASH. Genes analyzed in panels B and C are highlighted in red. Key components measured include: Tumor necrosis factor superfamily member 1A/2 (TNFR1/2); Interleukin-1 receptor (IL-1R); Toll-like receptor 4 (TLR4); Nuclear factor-kappa B (NF-κB); Mitogen-activated protein kinase (MAPK); Activator protein-1 (AP-1); Janus kinase (JAK); Signal transducer and activator of transcription (STAT); Tumor necrosis factor-alpha (*Tnfα*/TNFα); Interleukin 1-beta (*Il-1β*/IL-1β); Interleukin-6 (*Il-6*/IL-6); Transforming growth factor-beta (receptor) (TGF-β[R]); Suppressor of Mothers against Decapentaplegic 2/3/4 (SMAD2/3/4); Insulin-like growth factor-1 (receptor) (IGF-1[R]); *Collagen type I alpha I chain* (*Col1a1*); *Alpha-actin 2* (*Acta2*); Extracellular matrix (ECM); Matrix metalloproteinase 2/9 (MMP2/9); Tissue inhibitor of metalloproteinase 1/2 (TIMP1/2). **B,C)** Expression levels of inflammation- (B) and fibrosis-related (C) genes normalized to *Glyceraldehyde 3-phosphate dehydrogenase* (*Gapdh*) and shown as fold change relative to LF-fed SPF mice after 8 weeks determined via the 2^−ΔΔCt^ method. Data represent means ± SEM, analyzed via 3-way ANOVA (factors: Cholesterol, Fat, Time) followed by Tukey’s multiple comparisons. Bars with the same letter are not significantly different (*P*>0.05).

Gene expression in LX-2 cells (**Figure 5B,C**), TBA, and LPS in cecal homogenates (**Figure S5**) were analyzed via 3-way ANOVA following a 2 x 2 x 2 factorial design (Factors: Cholesterol, Fat, Microbes) in SPF and GF mice at 24 weeks. If a significant interaction or main effect was detected (*P*<0.05), posthoc Tukey’s multiple comparisons were performed.

Permutational multivariate analysis of variance (PERMANOVA) using ADONIS was performed on distance-based β-diversity matrices (Bray-Curtis, Weighted UniFrac, Unweighted UniFrac) to assess the main effects and interactions of time, cholesterol, and fat. Microbiome Multivariable Associations with Linear Models 2 (MaAsLin2)^37^ was performed using relative abundance feature tables obtained from QIIME2 to identify differential ASVs between treatment groups. The reference level was set to “LF”, and only significant (*P*<0.05) associations are shown.

Linear regression modeling of fecal amplicon sequences vs. fibrosis was performed using the nlme package (v 3.1-165) in R:

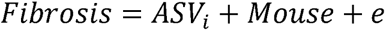

Where *Fibrosis* is the percent area stained with Picrosirius Red at 24 weeks, *ASV* is the relative abundance for (1-*i*) ASVs, *Mouse*; is the random effect of each animal, and *e* are random residual effects. An autoregressive variance-covariance matrix was applied to account for repeated measures within mice. Only ASVs that showed a significant effect (*P*<0.05) on fibrosis at each time point are shown.

## RESULTS

### SPF and GF mice show similar energy balance and body composition across diets over time

We first investigated how gut microbes interact with two prominent Western dietary components, cholesterol and saturated fat, in shaping MASLD/MASH outcomes. To test this, 16-week-old male GF and SPF C57Bl/6 mice were provided fructose- and glucose-supplemented drinking water and fed *ad libitum* one of four semi-purified diets: Low-fat (LF), Low-fat + Very High Cholesterol (LFVHC), High-fat (HF), or High-fat + Very High Cholesterol (HFVHC) (**Table S1**) for 24 weeks (**Figure 1A**).

Both SPF and GF mice fed HF or HFVHC exhibited increased percent body mass gain relative to baseline, demonstrating a significant main effect of dietary fat level and time (Fat *P*<0.001, Time *P*<0.001, respectively, **Figure 1B**). Percent body mass gain was evident throughout the study in both SPF HF and HFVHC-fed mice, whereas their GF counterparts showed a delay in weight gain, achieving significance after 4 weeks (**Figure 1B**). However, there was no overall effect of microbial status, suggesting that SPF and GF mice gain body mass similarly in response to dietary fat and cholesterol (**Figure 1B**). Total caloric intake mirrored these findings. In SPF mice, total caloric intake was influenced by an interaction between fat and time (Fat x Time *P*<0.001) as well as between cholesterol and fat (Cholesterol x Fat *P*=0.043), (**Figure S1A**). In GF mice, a significant main effect of cholesterol (Cholesterol *P*=0.028) and a fat x time interaction (Fat x Time *P*=0.030) was detected (**Figure S1A**. This indicated that dietary cholesterol alone contributed to increased caloric intake independent of saturated fat or time on diet. Similar to that observed for body mass gain, microbial status did not affect caloric intake throughout the study (**Figure 1B**). Together, these data suggest SPF and GF mice exhibit similar body weight gain and food consumption, regardless of dietary saturated fat or cholesterol level.

We next examined indicators of body composition, including liver and white adipose tissue mass. Liver mass, expressed as a percent of final body mass (LM%BM) was significantly impacted in SPF mice by an interaction between dietary cholesterol, fat, and time (Cholesterol x Fat x Time *P*=0.043, **Figure S1B**, left panel). At 8 weeks, SPF HFVHC diet-fed mice exhibited increased LM%BM relative to LF- and HF-fed counterparts, and this difference persisted relative to LF-fed mice at 24 weeks (**Figure S1B**, left panel). At 24 weeks, dietary fat alone elicited a significant main effect (Fat *P*<0.001), where both SPF and GF mice fed HF and HFVHC exhibited elevated LM%BM relative to LF and LFVHC groups (**Figure S1B**, right panel). These data suggest that diet-induced increases in LM%BM, a gross indication of MASLD, occurs in both GF and SPF mice, which becomes more pronounced over time.

Expansion of peripheral adipose tissue, another hallmark of MASLD, was evaluated by summing the masses of four major WAT depots (inguinal, retroperitoneal, mesenteric, and epididymal), which was expressed as a percent of final body mass (WAT%BM). In SPF animals, WAT%BM was significantly affected by fat (Fat *P*<0.001) as well as a significant interaction between cholesterol and time (Cholesterol x Time *P*=0.021). HFVHC-fed SPF mice exhibited increased WAT%BM relative to LF-fed counterparts at 8 weeks, but this difference diminished by 24 weeks when all diet groups converged (**Figure S1C**). After 24 weeks, only dietary fat elicited a significant main effect on WAT%BM when comparing both GF and SPF groups (Fat *P*<0.001), with patterns similar to those observed for liver mass (**Figure S1B, S1C** right panels).

Taken together, these data demonstrate that SPF and GF mice exhibited comparable caloric intake, body mass gain, and expansion of both liver and WAT in response to dietary fat and cholesterol intake. While some temporal differences were observed, particularly a delay in weight gain in GF mice, the overall patterns converged over time, indicating that gross metabolic responses to diet were independent of gut microbes.

### The presence of high fat and cholesterol-induced gut microbes are required for diet-induced fibrosing MASH

We next examined parameters of liver function and histological indicators of disease in SPF and GF mice. SPF mice fed HFVHC exhibited significantly elevated circulating Alanine transaminase (ALT) levels as early as 4 weeks relative to all other groups, which persisted through week 12 (Cholesterol x Fat x Time *P*=0.041; **Figure 1C**, left panel). Conversely, ALT elevation in HFVHC-fed GF mice was modest and delayed, reaching significance after 24 weeks compared to LF-fed GF counterparts (Cholesterol x Fat x Time *P*=0.025; **Figure 1C**, right panel).

Histology was assessed in H&E-stained liver sections via NAFLD Activity Score (NAS) which integrates scores of steatosis, lobular inflammation, and hepatocyte ballooning as previously described^30^. Representative images are shown in **Figure 1D**. In SPF mice, cholesterol elicited a significant main effect on NAS at 8 weeks (Cholesterol *P*=0.016), although group comparisons were not significantly different (**Figure 1E**, top and middle panel). After 24 weeks, NAS was increased across all groups in SPF mice (Time *P*<0.001), driven mainly by cholesterol (Cholesterol *P*=0.004; **Figure 1E**, top/middle panels). Lobular inflammation, a key NAS component, was also elevated by dietary cholesterol (Cholesterol *P*=0.016) at 8 weeks, which was further impacted by both fat and cholesterol after 24 weeks in SPF mice (Cholesterol x Fat *P*=0.024; **Figure 1F**, top/middle panels). Conversely, GF mice showed significantly lower total NAS (Microbe *P*=0.027) and lobular inflammation scores (Microbe *P*=0.004) compared to SPF mice at 24 weeks, regardless of diet (**Figure 1E, F**, bottom panels).

Oil-Red-O staining of liver sections revealed robust effects of dietary cholesterol on hepatic steatosis at both 8 (Cholesterol *P*<0.001) and 24 (Cholesterol *P*=0.008) weeks (**Figure S1D,E**). After 8 weeks, SPF mice fed both LFVHC and HFVHC showed significantly increased steatosis (**Figure S1E**, top panel). While differences between SPF groups diminished at 24 weeks, similar patterns were observed, with interactions between both cholesterol and time (Cholesterol x Time *P*<0.001) and fat and time (Fat x Time *P*=0.010) (**Figure S1E**, top/middle panels). Importantly, steatosis was significantly influenced by interactions between cholesterol and microbial status at 24 weeks (Microbe x Cholesterol *P*=0.020), with SPF mice showing greater lipid accumulation compared to their GF counterparts (**Figure S1E**, middle/bottom panel). Under GF conditions, no differences were observed between diet groups after 24 weeks (**Figure S1E**, bottom panel). These data suggest that SPF and GF mice were equally susceptible to diet-induced weight gain and hepatic lipid accumulation where cholesterol intake, regardless of fat level, increased NAS early in disease. However, only SPF mice fed HFVHC developed early, and progressively more severe clinical markers of liver damage as indicated by increased ALT levels over time, highlighting a microbe-dependent effect in driving disease progression to MASH.

We next assessed hepatic fibrosis, a hallmark of advanced MASH, under both GF and SPF conditions using histology analysis. Picrosirius red staining of liver sections revealed a robust increase in collagen deposition in HFVHC-fed SPF mice after 24 weeks, but not at 8 weeks (Cholesterol x Fat x Time *P*<0.001) compared to all other groups (**Figure 1G,H**). The ∼2-fold increase in percent area stained at 24 weeks in SPF HFVHC-fed mice was driven by a significant interaction between cholesterol and saturated fat (Cholesterol x Fat *P*=0.003). This effect was absent in GF mice, demonstrated by a microbe-dependent interaction (Cholesterol x Fat x Microbe *P*<0.001). (**Figure 1H**, middle/bottom panels). Collagen deposition determined via Masson’s Trichrome staining in liver sections was consistent with Picrosirius red staining, where SPF mice showed increased fibrosis after 24 weeks relative to 8 weeks (Cholesterol x Fat x Time *P*<0.001), while GF mice showed reduced fibrosis relative to SPF counterparts after 24 weeks (Cholesterol x Fat x Microbe *P*<0.001) (**Figure S1F,G**). Although quantification of both Picrosirius red and Masson’s Trichrome staining showed clear differences, fibrosis staging via pathological scoring was less definitive (**Figure 1I**). SPF mice showed more advanced fibrosis stages after 24 weeks compared to SPF mice at 8 weeks and GF counterparts at 24 weeks, although no consistent differences were apparent between HF and HFVHC groups under SPF conditions (**Figure 1I**).

Taken together, these data suggest dietary fat and cholesterol synergize to accelerate disease progression, which is dependent on the presence of gut microbes. This microbial-mediated effect manifests as substantial hepatic fibrosis only in HFVHC-fed SPF mice after 24 weeks.

### Dietary cholesterol promotes proinflammatory and fibrosis-related gene expression signatures in SPF mice

To explore the molecular response to high dietary cholesterol and saturated fat in SPF mice, we examined hepatic expression levels of key pro-inflammatory and fibrogenic genes. As outlined in **Figure 2A** (left panel), *Toll-like receptor 4* (*Tlr4*), a pattern recognition receptor that is implicated in MASLD/MASH^38^, as well as downstream pro-inflammatory cytokines, including *Tumor necrosis factor-alpha* (*Tnfα*), *Interleukin-1 beta* (*Il-1β*), and *Interleukin-6* (*Il-6*), have been shown to be upregulated in MASH patients^39^. Consistent with increased lobular inflammation (**Figure 1F**), expression of *Tlr4*, *Il-1β*, and *Tnfα* increased ∼2-3-fold in HFVHC-fed SPF mice relative to LF-fed controls after 24 weeks, with significant main effects of both time and cholesterol (Time *P*=0.010, <0.001, <0.001; Cholesterol *P*=0.023, 0.003, 0.004, respectively). However, *Il-6* levels were not significantly different (**Figure 2B**).

Given that proinflammatory cytokines drive HSC activation and fibrogenesis^40^, we next measured fibrosis-related gene expression. For example, as shown in **Figure 2A** (right panel), TNFα and IL-1β have been shown to increase the expression of Tissue inhibitor of metalloproteinase 1 (TIMP1)^41^, which, along with Matrix metalloproteinases (MMP), contribute to the regulation of extracellular matrix (ECM) remodeling in the liver (**Figure 2A**). We observed main effects of cholesterol, fat, and time (*P*=0.018, 0.011, <0.001 respectively) on *Timp1* expression, which was significantly increased in HFVHC-fed SPF mice after 24 weeks compared to all other groups (**Figure 2C**). A similar pattern was observed for *Timp2*, with main effects of cholesterol (*P*=0.032) and time (*P*=0.008). HFVHC-fed SPF mice showed significantly higher *Timp2* expression than all other groups except HF-fed SPF mice. Main effects of fat (*P*=0.030) and time (*P*=0.006) were evident for *Mmp2*, with pairwise comparisons showing that HFVHC-fed SPF mice at 24 weeks had higher expression than LF-fed mice after 24 weeks and all groups at week 8 (**Figure 2C**). In contrast, *Mmp9* expression was influenced only by time (*P*<0.001), with HFVHC-fed mice showing significantly higher levels at 24 weeks than LFVHC-, HF-, and HFVHC-fed mice at 8 weeks (**Figure 2C**).

Additionally, pro-fibrotic pathways in the liver are partly driven by Transforming growth factor beta (TGF-β), the protein product of *Tgfβ1*, which promotes expression of *Collagen type I alpha I chain* (*Col1a1*) and *Alpha-actin 2, smooth muscle* (*Acta2*). Conversely, Insulin-like growth factor-1 (IGF-1) is proposed to protect against hepatic fibrosis through several mechanisms (**Figure 2A**). We observed a time-dependent increase in *Tgfβ1* expression (*P*=0.016), though no significant pairwise differences were apparent (**Figure 2C**). Expression of *Igf1* was significantly impacted by main effects of saturated fat (*P*=0.032) and time (Time *P*<0.001), where LFVHC- and HF-fed mice at week 24 showed reduced *Igf1* expression relative to LF-fed mice at week 8 (**Figure 2C**). *Col1a1* expression was increased by dietary cholesterol, saturated fat, and time (*P*=0.021, 0.008, <0.001, respectively). HFVHC-fed mice at week 24 showed significantly higher *Col1a1* expression relative to LF-fed mice at week 24 as well as LF-, LFVHC-, and HF-fed mice at week 8 (**Figure 2C**). Acta2 expression was impacted only by time (*P*=0.001), with HFVHC-fed mice at week 24 exhibiting higher expression than LFVHC- and HF-fed mice at week 8 (**Figure 2C**).

These data suggest localized hepatic expression of growth factors in the liver involved in regulating fibrosis are influenced more by time and saturated fat than by dietary cholesterol. Specifically, *Tgfβ1* increases while *Igf1* decreases over time, with saturated fat contributing to an overall decrease in hepatic *Igf1* expression. However, it is important to consider that key regulators like TGF-β are also produced by nonparenchymal cells, such as immune cells and endothelial cells, which were not directly evaluated in this study^42^. These data, coupled with histological outcomes, suggest that dietary cholesterol-driven inflammation emerges early in SPF mice, while fibrosis develops over time and is further modulated by dietary saturated fat, which appears to be necessary, but not sufficient, on its own to drive fibrosis (**Figure 1G-I; S1F-G**).

### Gut microbiota diversity and composition are altered by dietary cholesterol and saturated fat during the development and progression of MASH

The composition of gut microbiota is rapidly altered in response to dietary modifications^43^, and our findings indicate their crucial role in triggering diet-induced fibrosing MASH. To examine the impact of dietary cholesterol vs. saturated fat on gut microbiota composition, we performed 16S rRNA gene amplicon sequencing. To ascertain a dose-dependent effect of dietary cholesterol, we included two additional diets: low-fat high-cholesterol (LFHC) and high-fat high-cholesterol (HFHC), each of which contain 0.2% cholesterol (**Table S1**).

We first assessed overall microbial community membership via α-diversity (within-sample diversity) metrics in feces collected every 4 weeks. Reductions in richness (Chao1) and richness/evenness (Fisher’s alpha) were evident as early as 4 weeks on HFVHC diet (Cholesterol x Fat x Time *P*<0.001 for both; **Figure S2A**). Both Shannon index, another richness/evenness metric, and Simpson index, a measure of species dominance, were altered by week 8 (Time *P*<0.001, Cholesterol x Fat *P*<0.001 for both; **Figure S2A**).

Examination of α-diversity in cecal contents collected after 8 and 24 weeks showed that Chao1 richness was reduced in response to diet, driven by an interaction between cholesterol and saturated fat (Cholesterol x Fat *P*=0.048), demonstrating a consistent trend toward reduced richness in HFVHC-fed mice (**Figure S2B**). A similar trend was observed in Fisher’s alpha, though statistical significance was not achieved (**Figure S2B**). Shannon index was affected by both time and a cholesterol x fat interaction (Time *P*<0.001, Cholesterol x Fat *P*=0.031), while Simpson index was increased, indicating lower diversity, due to main effects of saturated fat and cholesterol (Fat *P*=0.001, Cholesterol *P*=0.010; **Figure S2B**). The latter findings contrast with those observed in feces (**Figure S2A**), suggesting regional differences in HFVHC-induced changes to α-diversity.

We next investigated between-sample β-diversity metrics to determine how diet alters microbial community membership across time. In feces, nonphylogenetic-based Bray-Curtis analysis showed significant interactions between cholesterol and saturated fat (Cholesterol x Fat, *P*=0.001), cholesterol and time (Cholesterol x Time, *P*=0.003), and saturated fat and time (Fat x Time *P*=0.002), indicating that these dietary components dynamically shift fecal microbiota composition over time (**Figure S2C**). These interactions were evident at week 4 (*P*=0.001), and persisted at all subsequent time points, including week 8 (*P*=0.004), 12 (*P*=0.001), 16 (*P*=0.003), 20 (*P*=0.045), and 24 (*P*=0.019) (**Figure S2D**). Importantly, HFVHC-fed mice demonstrated a pronounced and persistent separation from all other groups beginning at week 8, revealing that the combination of saturated fat and cholesterol drove a distinct microbial composition throughout the study (**Figure S2D**).

To determine the regional influence of saturated fat vs. cholesterol over time, we examined β-diversity of cecal microbes at 8 and 24 weeks (**Figure 3A; S2E,F**). At both time points, ADONIS analysis showed microbiota community membership was significantly altered by interactions between saturated fat and cholesterol, as indicated by shifts in both non-phylogenetic-based (Bray-Curtis: 8 weeks *P*=0.004; 24 weeks *P*=0.002; **Figure 3A**) and phylogenetic-based (Unweighted: 8 weeks *P*=0.018; 24 weeks *P*=0.001; Weighted: 8 weeks *P*=0.007; 24 weeks *P*=0.021; **Figure S2E,F**) distance metrics.

**Figure 3.**
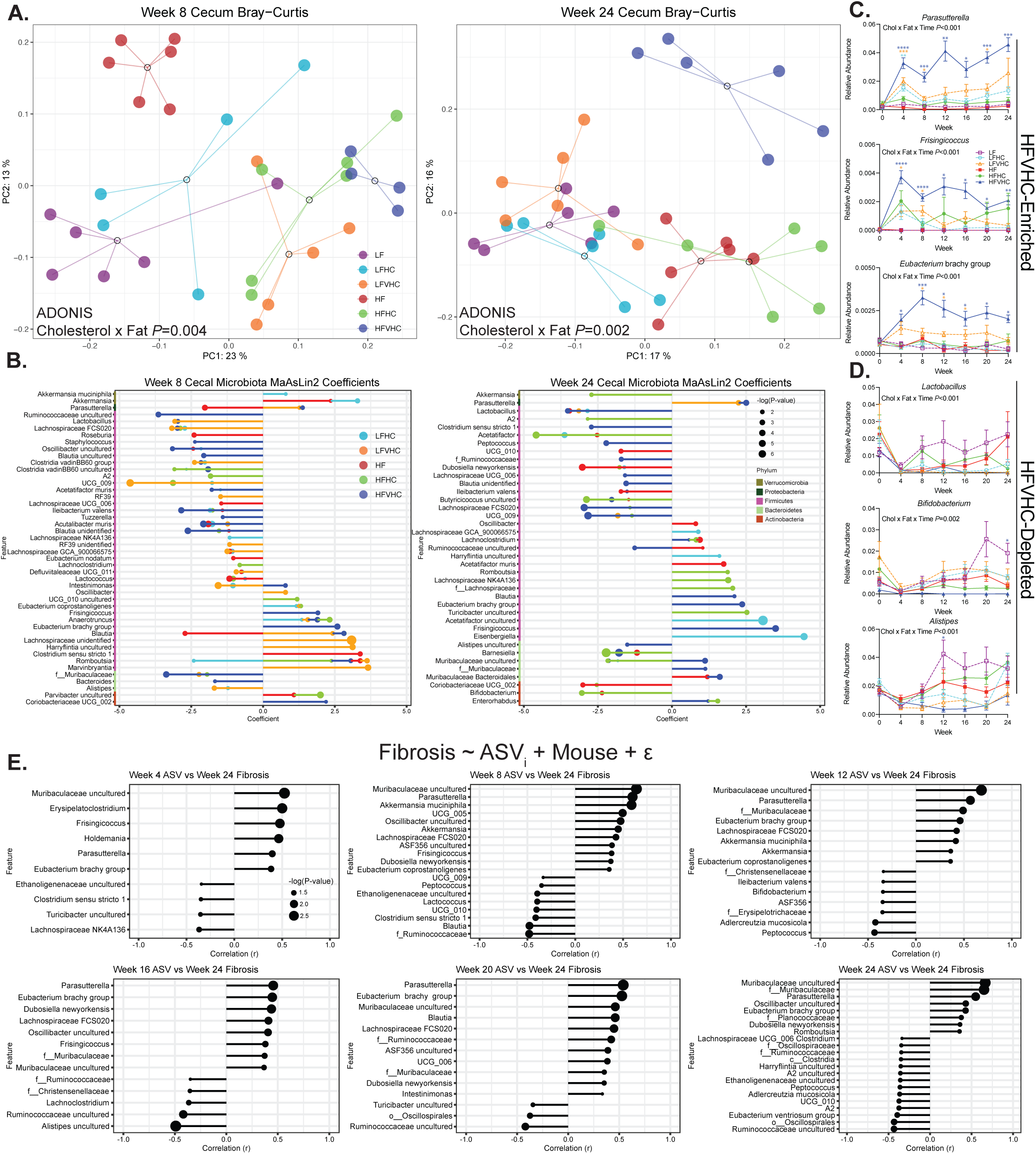
Dietary cholesterol and saturated fat lead to rapid and sustained shifts in gut microbiota that are associated with hepatic fibrosis. **A)** Bray-Curtis β-diversity PCoA of cecal microbiota after 8 (left panel) and 24 (right panel) weeks. Dots represent individual mice, open circles represent centroids with lines connecting individual dots within a treatment group. Data analyzed via multifactorial ADONIS (Factors: Cholesterol, Fat). **B)** ASVs significantly enriched or depleted relative to LF-fed mice via multivariate MaAsLin2 analysis. Positive coefficients indicate enrichment; negative coefficients indicate depletion. **C,D)** Relative abundance of fecal ASVs enriched (C) or depleted (D) in response to HFVHC over time. Data represent means ± SEM, analyzed via 3-way ANOVA (Factors: Cholesterol, Fat, Time) followed by Tukey’s multiple comparisons within timepoint. \**P*<0.05, \*\**P*<0.01, \*\*\**P*<0.005, \*\*\*\**P*<0.001. Asterisk color represents significance relative to LF-fed mice. **E)** Linear modeling of relationships between fecal ASVs in feces collected every four weeks throughout the study relative to severity of hepatic fibrosis (based on Picrosirius red-stained area) determined at 24 weeks. r, Pearson correlation coefficient; dot size corresponds to -log(P-value). For B and E, Dot size reflects -log(P-value). Taxonomic labels: “f__”: family-level annotation; “o__” order-level annotation; “c__” class-level annotation.

To identify specific ASVs that contribute to the observed shifts in microbial composition, we performed Microbiome Multivariable Associations with Linear Models 2 (MaAsLin2)^37^. After 8 weeks, several ASVs were enriched in HFVHC-fed mice, including *Parasutterella*, *Instestimonas*, *Frisingicoccus*, *Anaerotruncus*, *Eubacterium* brachy group, *Blautia*, *Romboutsia*, and *Coriobacteriaceae* UCG_002 (**Figure 3B**, left panel). Conversely, LF-fed mice showed enrichment of taxa such as Ruminococcaceae uncultured, *Lactobacillus*, *Lachnospiraceae* FCS020, *Staphylococcus*, *Oscillobacter* uncultured, *Blautia* uncultured, *Clostridia* vandin BB60 uncultured, *Acetatifactor muris*, *Ileibacterium valens*, *Tuzzerella*, *Acutalibacter muris*, *Blautia* unidentified, *Muribaculaceae*, and *Bacteroides* (**Figure 3B**, left panel).

Interestingly, several ASVs enriched by HFVHC diet, including *Parasutterella*, *Frisingicoccus*, *Eubacterium* brachy group, and *Blautia*, remained elevated after 24 weeks, while *Lachnoclostridium*, Muribaculaceae, and *Enterorhabdus* emerged only at this later timepoint (**Figure 3B**, right panel). LF-fed mice continued to show enrichment of Ruminococcaceae uncultured, *Lactobacillus*, and *Lachnospiraceae* FCS020, whereas additional taxa, including *Clostridium* sensu stricto 1, *Peptococcus*, Ruminococcaceae, *Lachnospiraceae* UCG_006, *Blautia* unidentified, *Ileibacterium valens*, *Butyricoccos* uncultured, UCG_009, *Alistipes* uncultured, and *Barnesiella*, were exclusively enriched at 24 weeks (**Figure 3B**, right panel).

Pairwise comparisons of cecal ASVs between diets at 8 and 24 weeks via MaAsLin2 revealed differential and time-dependent effects of cholesterol vs. saturated fat on microbial community membership. The addition of very high cholesterol to the HF diet dramatically impacted ASV relative abundances at both timepoints (**Figure S3A,B**), consistent with our findings related to β-diversity (**Figure 3A; S2C-E**). Importantly, *Parasutterella* was the only ASV consistently enriched in response to high dietary cholesterol, regardless of saturated fat level or timepoint (**Figure S3A-D**). Further, *Bifidobacterium* was reduced in HFVHC-fed mice relative to HF only after 24 weeks (**Figure S3B**), indicating a delayed response to dietary cholesterol (**Figure S3A**). Conversely, *Lactobacillus* was reduced in LFVHC-fed mice relative to LF at both timepoints, however, no differences were observed between HF and HFVHC-fed mice. This may suggest high dietary saturated fat intake masks the influence of cholesterol on *Lactobacillus* abundance over time (**Figure S3A,B**). When examining the effects of saturated fat alone, i.e., in the absence of added cholesterol, more pronounced shifts in cecal microbial composition were evident at 8 weeks than at 24 weeks (**Figure S3C,D**). Although few taxa showed consistent patterns at both timepoints, *Ileibacterium valens* was reduced at 8 and 24 weeks, whereas *Bifidobacterium* and *Lactobacillus* were both reduced in response to saturated fat after 24 weeks (**Figure S3D**).

To better understand how these microbial shifts identified via MaAsLin2 evolved throughout the course of disease, we examined the relative abundances of key microbial taxa in monthly fecal samples (**Figure 3C**). This analysis revealed that many diet-induced shifts were observed as early as 4 weeks. HFVHC diet enriched *Parasutterella*, *Frisingicoccus*, and *Eubacterium* brachy group, which persisted throughout the study (Cholesterol x Fat x Time *P*<0.001 for all; **Figure 3C**). Other taxa, such as *Lactobacillus*, *Bifidobacterium*, and *Alistipes,* were robustly depleted in HFVHC-fed mice relative to LF-fed controls (Cholesterol x Fat x Time *P*<0.001, 0.002, <0.001, respectively; **Figure 3D**).

Finally, linear modeling was applied to fecal ASVs across time at 4, 8, 12, 16, 20, and 24 weeks to identify taxa that significantly correlated with fibrosis severity based on Picrosirius red staining at 24 weeks (**Figure 1E**). Notably, several taxa identified by MaAsLin2 as enriched in HFVHC-fed mice, including *Parasutterella*, *Frisingicoccus*, *Eubacterium* brachy group, and members of *Muribaculaceae*, also showed an early and sustained association with hepatic fibrosis, suggesting these microbes may act as disease drivers (**Figure 3E**).

These microbial shifts raise the question of how dietary interventions may impact host-microbiota metabolic interactions, particularly bile acid (BA) metabolism, which plays a central role in gut-liver axis regulation. Notably, *Parasutterella*—a genus enriched early in HFVHC-fed mice and associated with more severe fibrosis—is a member of the Proteobacteria phylum, which tends to exhibit high tolerance to BAs^44^. The introduction of *Parasutterella* to a complex gut microbial community leads to decreases in cholic acid (CA), taurocholic acid (TCA), taurodeoxycholic acid (TDCA), 7-ketoDCA, and glycolithocholic acid (GLCA) sulfate in the cecum, suggesting it may either possess BA metabolization capabilities and/or alter the abundance/activity of other BA metabolizing microbiota^45^. Conversely, *Bifidobacteria* and *Lactobacillus* are more sensitive to high concentrations of BAs^44^. Further, many members of these genera have well-established capabilities to perform deconjugatation and transformation of BAs in the gut^46,47^. Given the complex interplay between microbial composition, bile acid dynamics, and liver health, we next investigated how dietary cholesterol and saturated fat shape BA metabolism over time.

### High dietary cholesterol and saturated fat alter fecal bile acid composition over time in distinct and synergistic ways in SPF mice

BAs, synthesized in the liver from cholesterol, are a key component in lipid digestion and absorption and play a central role in maintaining the gut-liver axis through their signaling functions and interactions with gut microbes. Given the significant shifts observed in gut microbes in SPF mice, we examined how dietary cholesterol and saturated fat affect host hepatic expression of genes involved in BA metabolism. We first assessed hepatic expression of genes that encode key enzymes involved in host BA synthesis, mainly *Cytochrome P450 family 7 subfamily A member 1* (*Cyp7a1*), which catalyzes the rate-limiting step in the classic BA biosynthetic pathway, as well as *Cytochrome P450 family 27 subfamily A member 1* (*Cyp27a1*), which is associated with the alternative BA biosynthetic pathway. We noted that dietary cholesterol and time significantly impacted *Cyp7a1* expression (Cholesterol *P*=0.005; Time *P*=0.002) (**Figure 4A**, top panel). LFVHC- and HFVHC-fed mice exhibited numerically elevated expression of *Cyp7a1* at 8 weeks, although these increases did not reach statistical significance at this timepoint (**Figure 4A**, top panel). Interestingly, *Cyp7a1* in LFVHC-fed mice at 8 weeks was significantly increased relative to LF, LFVHC, and HF-fed groups at week 24; however, no differences were observed among groups within the 24 week timepoint (**Figure 4A**, top panel), suggesting a transient impact of dietary cholesterol on BA synthesis genes. No significant differences were observed in *Cyp27a1* gene expression across all groups or timepoints, indicating a limited role for the alternative pathway in this context (**Figure 4A**, bottom panel). This suggests high dietary cholesterol intake exerts early effects on host BA metabolism mediated by the classic biosynthesis pathway, which are lost following long-term exposure.

**Figure 4.**
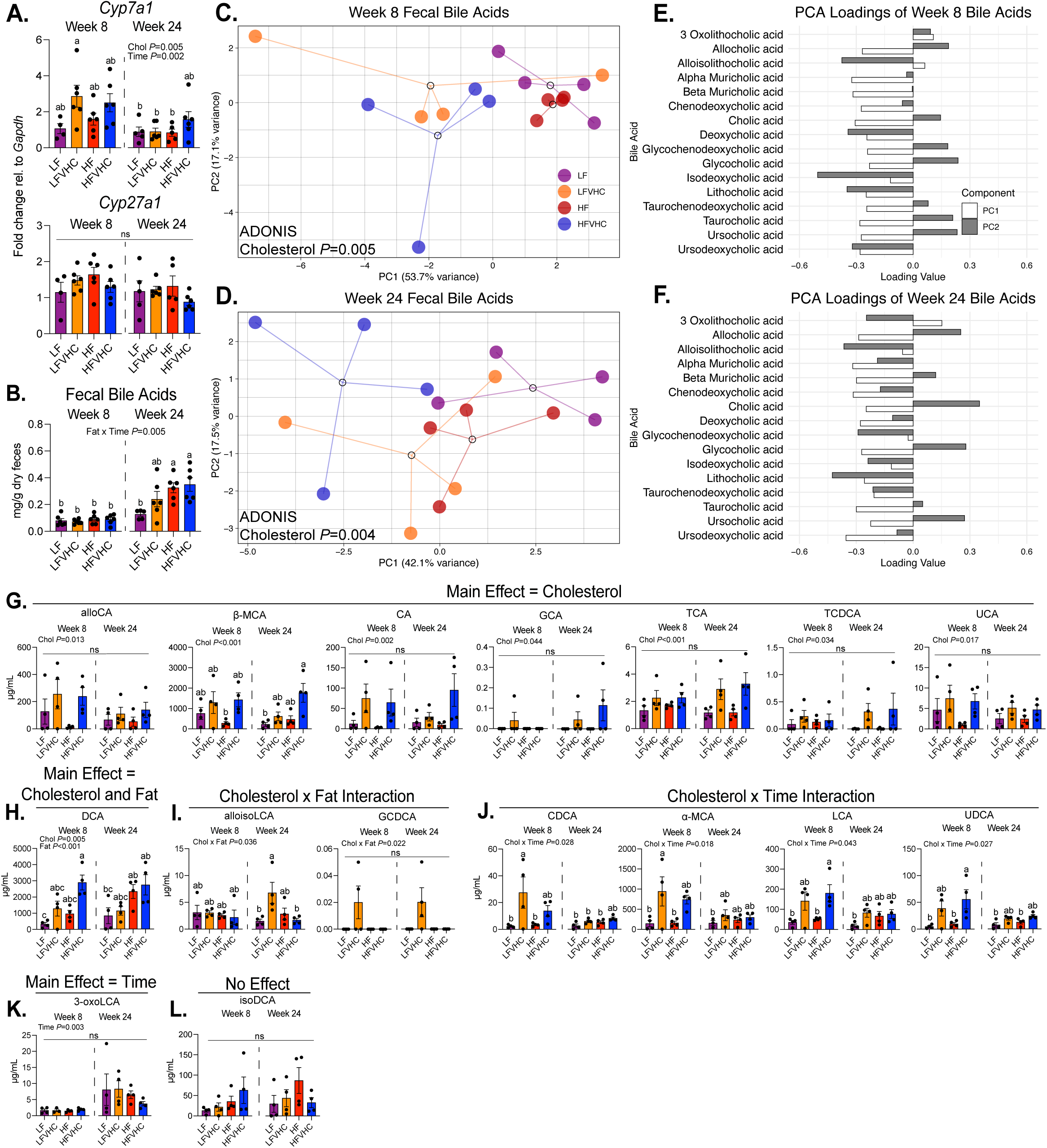
Time- and diet-dependent remodeling of fecal bile acid composition by cholesterol and saturated fat in SPF mice. **A)** Hepatic expression of genes involved in BA biosynthesis shown as fold change relative to LF-fed SPF mice at 8 weeks, normalized to *Gapdh* and determined via the 2^−ΔΔCt^ method. *Cytochrome P450 family 7 subfamily A member 1* (*Cyp7a1*, top panel); *Cytochrome P450 family 27 subfamily A member 1* (*Cyp27a1*, bottom panel). **B)** Fecal BA concentrations in SPF mice. Data represent means ± SEM, analyzed via 3-way ANOVA (factors: Cholesterol, Fat, Time) followed by Tukey’s multiple comparisons. Bars with the same letter are not significantly different (*P*>0.05). **C,D)** PCA plots of fecal BA from SPF mice at 8 weeks (C) and 24 weeks (D). Dots represent individual mice, open circles represent centroids with lines connecting individual dots, analyzed via multifactorial ADONIS (Factors: Cholesterol, Fat). **E,F)** PC loadings of individual BAs after 8 weeks (E) and 24 weeks (F). **G-L)** Quantification of BA in feces of SPF mice at 8 and 24 weeks significantly impacted by a main effect of dietary cholesterol (G), main effects of dietary cholesterol and saturated fat (H), interaction between dietary cholesterol and saturated fat (I), interaction between dietary cholesterol and time (J), a main effect of time (K), or not impacted by any factor (L). Data represent means ± SEM, analyzed via 3-way ANOVA (factors: Cholesterol, Fat, Time) followed by Tukey’s multiple comparisons. Bars with the same letter are not significantly different (*P*>0.05).

We next examined total fecal BA concentrations and individual BA species. An interaction between fat and time (Fat x Time *P*=0.005) elicited a significant impact on total fecal BA levels, where HF and HFVHC-fed mice at 24 weeks exhibited higher concentrations compared to all other groups, except LFVHC at 24 weeks, a group that showed intermediate levels (**Figure 4B**). These data suggest that while cholesterol influences BA synthesis, saturated fat level may alter BA excretion and composition over time.

Lipidomic analysis of fecal BAs was performed in a subset of SPF mice at 8 and 24 weeks showing that of the 40 identified BAs, 16 were quantifiable with distinct patterns influenced by dietary cholesterol and fat. A heatmap of all 40 detected BAs (including both primary and secondary as well as conjugated and unconjugated species) is shown in **Figure S4A**. Principal Component Analysis (PCA) of the 16 quantified BAs revealed that dietary cholesterol significantly altered BA profiles at both 8 (*P*=0.005) and 24 weeks (*P*=0.004) (**Figure 4C,D**). PC loading plots were assessed to identify specific BAs driving these compositional shifts (**Figure 4E,F**). At 8 weeks, α-Muricholic acid (α-MCA), β-Muricholic acid (β-MCA), chenodeoxycholic acid (CDCA), deoxycholic acid (DCA), isoDCA, lithocholic acid (LCA), and ursodeoxycholic acid (UDCA) exhibited negative loadings on both PC1 and PC2 (**Figure 4E**), indicating stronger associations with HFVHC-fed mice, contributing to the separation of this group in the ordination plot (**Figure 4C**). At 24 weeks, allocholic acid (alloCA), β-MCA, CA, glycocholic acid (GCA), taurocholic acid (TCA), and ursocholic acid (UCA) showed similar loading outcomes (**Figure 4D,F**).

Absolute quantifications of these BAs in feces are shown in **Figure 4G-L**. While saturated fat alone did not elicit a significant main effect on any BA, a significant main effect of cholesterol was observed for several species, including alloCA (*P*=0.013), β-MCA (*P*<0.001), CA (*P*=0.002), GCA (*P*=0.004), TCA (*P*<0.001), and taurochenodeoxycholic acid (TCDCA) (*P*=0.034) (**Figure 4G**). Fecal β-MCA concentration was higher in HFVHC-fed mice compared to both LF-fed mice after 24 weeks and HF-fed mice after 8 weeks (**Figure 4G**). DCA levels were significantly impacted by both cholesterol (*P*=0.005) and fat (*P*<0.001), where HFVHC-fed mice exhibited the highest fecal DCA concentrations relative to LF-fed mice at both 8 and 24 weeks (**Figure 4H**). Further, significant interactions between cholesterol and fat were evident for both alloisoLCA (*P*=0.036) and glycochenodeoxycholic acid (GCDCA) (*P*=0.022) (**Figure 4I**). Pairwise comparisons showed that LFVHC-fed mice had significantly higher alloisoCA compared to both LF- and HFVHC-fed counterparts at 24 weeks (**Figure 4H**). Other BAs showed significant interactions between cholesterol and time. For instance, α-MCA was significantly impacted (*P*=0.018), where LFVHC-fed mice at 8 weeks showed higher levels than LF- and HF-fed mice at 8 weeks, as well as LF-fed mice at 24 weeks (**Figure 4J**). Likewise, CDCA (*P*=0.028) was increased in LFVHC-fed mice at 8 weeks compared to all groups except HFVHC-fed mice at both 8 and 24 weeks (**Figure 4J**). Both LCA (*P*=0.043), and UDCA (*P*=0.027) showed similar trends, with the highest concentrations observed in mice fed HFVHC at 8 weeks (**Figure 4J**). 3-oxoLCA was increased over time, regardless of diet (Time *P*=0.003; **Figure 4K**), while isoDCA was not differentially impacted by factors of diet or time (**Figure 4L**).

Together, these data show that prolonged intake of high dietary cholesterol and saturated fat synergistically remodel the BA pool in SPF mice. Given the well-established link between BAs, gut microbes, and hepatic inflammation and fibrosis, these shifts may contribute to the fibrotic phenotype observed only in SPF mice after prolonged HFVHC feeding.

### Gut microbiota-dependent components modulated by HFVHC diet activate human hepatic stellate cells *in vitro*

Given the observed shifts in gut microbiota and BAs in response to high levels of dietary cholesterol and saturated fat that coincided with the development of fibrosing MASH, we next tested whether gut-derived factors induced by HFVHC feeding could directly drive fibrogenic processes. To address this, cell-free cecal homogenates, i.e., cecal water, were generated from GF and SPF mice fed LF, LFVHC, HF, or HFVHC diet for 24 weeks. These homogenates were applied to human-derived LX-2 HSCs *in vitro*, as outlined in **Figure 5A**. After 4 hours of exposure, expression of both pro-fibrogenic and pro-inflammatory genes were measured via qRT-PCR. We observed a significant three-way interaction between dietary cholesterol, saturated fat, and microbial status for cecal homogenates on *COL1A1* (*P*=0.025) and *TGFβR2* (*P*=0.015) (**Figure 5B**). Cecal homogenates from HFVHC-fed SPF mice drove a significant upregulation of *COL1A1* and *TGFβR2* expression ∼10 and ∼4-fold, respectively, relative to all other groups (**Figure 5B**).

**Figure 5.**
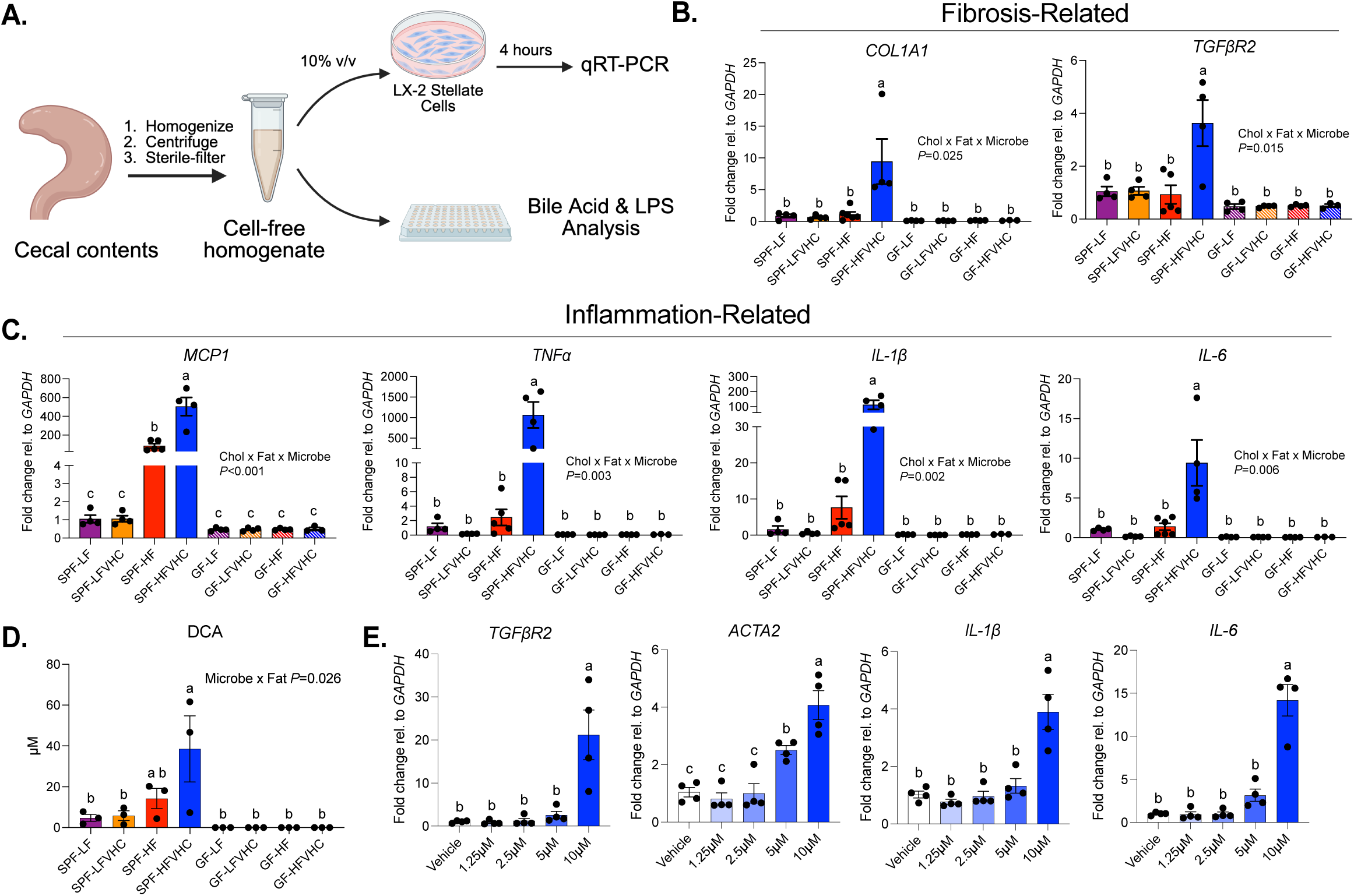
Microbiota-dependent gut factors from HFVHC-fed SPF mice induce pro-fibrotic and pro-inflammatory activation of hepatic stellate cells *in vitro*. **A)** Experimental schematic; cecal contents were from pooled from 3 mice/group after 24 weeks on diet, followed by homogenization, centrifugation, and sterile filtration, followed by TBA and LPS quantification. LX-2 cells were exposed to homogenates at 10% media (v/v) for 4 hours followed by qRT-PCR. Created on Biorender.com. **B,C)** Gene expression levels of fibrosis-related (B) and inflammation-related (C) HSC activation markers. *Collagen type I alpha I chain* (*COL1A1*), *Monocyte chemoattractant protein-1* (*MCP1*), *Interleukin-6* (*IL-6*), *Transforming growth factor-beta receptor type 2* (*TGFβR2*), *Tumor necrosis factor-alpha* (*TNFα*), *Interleukin 1-beta* (*IL-1β*). **D)** Concentration of DCA in cecal homogenates. **E)** LX-2 cells were treated with indicated concentrations of DCA or vehicle for 4 hours followed by qRT-PCR. Alpha-smooth muscle actin (*ACTA2*). Data represent means ± SEM. (B-D) analyzed via 3-way ANOVA (factors: Time, Fat, Microbe) followed by Tukey’s multiple comparisons within timepoint. (E) analyzed via 1-way ANOVA followed by Tukey’s multiple comparisons. Bars with the same letter are not significantly different (*P*>0.05). Gene expression values (C,E) normalized to *GAPDH* and determined via the 2^−ΔΔCt^ method and expressed relative to SPF-LF (B,C) or Vehicle (E) groups.

We next investigated the expression of pro-inflammatory markers in LX-2 HSCs following exposure to cecal homogenates. Similar to what was observed for pro-fibrotic genes, a significant three-way interaction between dietary cholesterol, saturated fat, and microbial status was observed for *Monocyte chemoattractant protein-1* (*MCP1*) (*P*<0.001), *TNFα* (*P*=0.003), *IL-1β* (*P*=0.002), and *IL-6* (*P*=0.006) (**Figure 5C**). Pairwise comparisons showed that cecal homogenates from HFVHC-fed SPF mice significantly increased expression of all four activation markers relative to all other groups. Here, *MCP1*, *TNFα*, and *IL-1β* were robustly upregulated (∼500-, 1200-, and 100-fold, respectively) in LX-2 HSCs exposed to HFVHC cecal homogenate, while *IL-6* showed a 10-fold increase (**Figure 5C**). Interestingly, *MCP1* expression was also significantly increased following exposure to cecal homogenate from HF-fed SPF mice relative to all groups except SPF-HFVHC (**Figure 5C**). These data suggest that HFVHC feeding induces gut factors that elicit strong activation of pro-fibrogenic and inflammatory pathways in HSCs *in vitro*, providing a mechanistic link between diet, gut microbes and hepatic injury.

To gain further insights into microbially-derived factors in HFVHC cecal homogenates that may drive upregulation of pro-fibrotic and pro-inflammatory genes in LX-2 HSCs, we examined lipopolysaccharide (LPS) and total bile acid (TBA) concentrations^48,49^. LPS levels were impacted by multiple two-way interactions, including cholesterol x microbial status (*P*<0.001), cholesterol x fat (*P*<0.001), and fat x microbial status (*P*=0.001). While HFVHC-fed SPF cecal homogenates had higher LPS levels compared to LF-fed SPF and HFVHC-fed GF groups, no significant differences were apparent compared to other groups (**Figure S5B**). TBA concentrations exhibited a significant three-way interaction between dietary cholesterol, fat, and microbial status (*P*<0.001, **Figure S5A**). Surprisingly, cecal homogenate from LFVHC-fed SPF mice had the highest TBA levels, followed by homogenate from LF-fed SPF mice, both of which were significantly greater than all other groups (**Figure S5A**). These results suggest that neither LPS nor TBA concentration alone in cecal homogenates is not the main driver of HSC activation observed *in vitro*, but specific BAs may be enriched in HFVHC-fed SPF mice.

Analysis of BA species in cecal homogenates revealed an interaction between microbial status and fat (*P*=0.026) for DCA (**Figure 5D**), where HFVHC-fed SPF mice exhibited significantly increased levels relative to all groups except for HF, although a numerical trend was evident. Interestingly, a similar trend was observed for ω/α-MCA, which also showed a significant interaction between microbial status and fat (*P*=0.019; **Figure S5C**). While both GCA and TCDCA were also impacted by a significant interaction between fat and microbial status (*P*<0.001 and *P*=0.002, respectively), cecal homogenate from SPF LF and LFVHC-fed mice showed higher levels relative to all other groups (**Figure S5D**). CA was impacted by both an interaction between microbial status and fat (*P*<0.001) as well as between microbial status and cholesterol (*P*=0.032) (**Figure S5E**). This BA showed similar trends as CDCA (*P*=0.015) and β-MCA (*P*=0.008), which also showed a significant interaction between microbial status and cholesterol, where cecal homogenate from LFVHC-fed mice showed the highest levels (**Figure S5F**). Both LCA and UDCA were impacted by a main effect of microbes (*P*=0.002 and *P*<0.001, respectively), where SPF mice showed higher levels than GF mice regardless of diet (**Figure S5G**).

Given that DCA showed the most distinct pattern of significance in cecal homogenate from HFVHC-fed SPF mice, we next tested whether this BA could activate HSCs *in vitro*. LX-2 cells were exposed to varying concentrations for 4 hours. We observed significantly increased expression in both fibrosis-related genes (*TGFβR2* and *ACTA2*) and inflammation-related genes (*IL-1β* and *IL-6*) in response to 10 µM DCA relative to vehicle control (**Figure 5E**). ACTA2 expression was also modestly increased by 5 µM DCA (**Figure 5E**).

Taken together, these data suggest that neither TBA nor LPS levels alone accounted for the observed HSC activation induced by cecal homogenates from HFVHC-fed SPF mice. However, fat and cholesterol-induced microbially-derived BA species, such as DCA, may serve as key drivers of the initial pro-inflammatory and fibrogenic activation of HSCs *in vitro.* Further work is needed to determine if DCA acts alone or in concert with additional BAs or other microbially-derived factors to activate HSCs. In addition, whether the sustained expansion of dietary cholesterol and fat-induced microbial community members, such as *Parasutterella*, directly contributes to increased DCA over time requires further exploration in the context of fibrosis in MASH.

## DISCUSSION

In humans, fibrosing MASH develops through a heterogeneous and dynamic process, where multiple factors, including diet and gut microbiota, have been established as robust disease moderators^8^. Yet, the specific interactions between common Western dietary components and gut microbiota imbalances in the context of MASLD to MASH progression are not fully understood^50–54^. Dietary cholesterol and saturated fat are often consumed together in human diets, making it difficult to disentangle their individual effects on gut microbiota and disease progression. By independently manipulating these two prominent Western dietary components, our study reveals their distinct and synergistic effects on gut microbiota composition and development of fibrosing MASH. Using a multifactorial design that incorporated dietary composition, microbial status (GF vs. SPF), and time, we were able to dissect both the individual and combined contributions of these variables to disease etiology in a well-established mouse model. Our findings support the notion that cholesterol and saturated fat remodel gut microbiota early in disease, leading to persistent alterations that contribute to fibrosing MASH over time.

We showed significant induction of hepatic fibrosis in SPF mice fed HFVHC diet compared to all other groups (**Figure 1G,H; Figure S1F,G**), suggesting dietary saturated fat, cholesterol, and the presence of gut microbes are necessary for the development of MASH with fibrosis *in vivo*. This finding was further supported via the robust activation of LX-2 HSCs *in vitro* by cecal homogenate from HFVHC-fed SPF mice (**Figure 5**). This suggests that gut-derived factors induced by the synergistic effects of cholesterol and fat may directly promote hepatic fibrogenesis through HSC activation, which could be mediated by several mechanisms^19,55–58^. Previous studies have shown that diets high in cholesterol (2% wt/wt) and fat (40% kcal) can increase the relative abundance of *Blautia producta,* a gut bacteria that produces 2-oleoylglycerol and activates hepatic macrophages in mice^59^. A separate study showed that 2% wt/wt inclusion of cholesterol shifted gut microbiota and BA profiles, including increased hepatic DCA and CDCA, which were sufficient to activate inflammatory gene expression in HepG2 cells^57^. To our knowledge, no study has directly investigated the effect of microbiota- and diet-dependent gut factors that stimulate HSCs. Additional studies are needed to identify the factor(s) responsible for HSC activation and to test their sufficiency both *in vitro* and *in vivo*. Further, given the complexity of the *in vivo* hepatic microenvironment, future validation in primary or human HSCs as well as *in vivo* models is warranted to confirm the pro-fibrotic role of DCA and other gut-derived factors.

Several gut microbial taxa exhibited early and sustained enrichment in response to HFVHC diet and were positively associated with hepatic fibrosis at multiple time points (**Figure 3B,C,D**). *Parasutterella* was particularly sensitive to cholesterol and showed a strong association with hepatic fibrosis (**Figure 3D**). Colonization of SPF mice with *Parasutterella* mc1 has been shown to alter gut BA profiles and reduce the gene expression of several ileal BA transporters and Farnesoid X Receptor (FXR) signaling pathway components, suggesting a role in regulating enterohepatic BA circulation^45^. Additionally, during antibiotic-induced dysbiosis, *Parasutterella excrementihominis* upregulates fatty acid biosynthesis pathways in the small intestine, highlighting its metabolic adaptability and potential to thrive under disrupted microbial conditions^60^. Despite the prevalence of this genus in humans and mice^61,62^, it remains relatively unexplored. Some studies have linked *Parasutterella* with health benefits, including reduced hypothalamic inflammation and improved LDL levels^61,63^. In contrast, other studies have associated increased *Parasutterella* abundance with hepatic steatosis^64^, type 2 diabetes^65^, and obesity^66^. In our study, *Parasutterella* was consistently enriched in mice with more severe hepatic fibrosis and rapidly expanded upon exposure to HFVHC feeding, prior to detectable inflammation or fibrosis. It is likely that *Parasutterella* exhibits strain-level heterogeneity that may explain its differential association with metabolic diseases. While *Parasutterella* was strongly associated with fibrosis and rapidly expanded in HFVHC-fed mice, further studies are required to determine causality in MASH.

In addition to direct interactions with sterols in the gut, both dietary cholesterol and saturated fat significantly impact BA metabolism and secretion^67^. BAs, in turn, can exert both beneficial and detrimental effects on the gut microbiota community and may represent a key mechanistic link between diet, shifts in gut microbiota, and fibrosing MASH. We observed significant alterations in the abundance of several BAs in feces, predominantly driven by dietary cholesterol, even in the absence of saturated fat (**Figure 4, S5**). DCA, a strongly hydrophobic and cytotoxic secondary BA^68^ produced by microbial dehydroxylation of CA, tended to be the highest in HFVHC-fed mice at both 8 and 24 weeks, while CA was highest in LFVHC-fed mice after 8 weeks and HFVHC-fed mice at both timepoints (**Figure 4G,H**). These findings suggest that increased levels of DCA are not solely dependent on CA availability, but rather on shifts in microbial capacity to metabolize CA.

These changes in BAs likely contribute to the depletion of certain microbial taxa. In our model, HFVHC feeding depleted *Lactobacillus*, *Bifidobacteria*, *Alistipes*, and other taxa negatively associated with hepatic fibrosis (**Figure 3B-D**). This trend is consistent with previous evidence suggesting that several members of *Bifidobacteria* and *Lactobacillus* are beneficial to human health^69^ and protect against liver disease and damage^70–78^. The depletion of these bacteria could be due to the observed shifts in BAs, particularly unconjugated BAs^79^, that negatively impact several members of *Bifidobacteria* and *Lactobacillus*. In our study, fecal DCA and CA concentrations tended to be the highest in HFVHC-fed SPF mice (**Figure 4G,H**), suggesting they may be partially responsible for the loss of BA-sensitive taxa. For instance, the growth of several *Bifidobacteria* species, including *B. longum*, *B. pseudolongum*, *B. adolescentis,* and *B. pseudocatenulatum*, was inhibited by CDCA, DCA, and CA *in vitro*^44^.

While BAs can differentially influence certain microbes, elevated DCA may also act directly on the liver to promote fibrosis. In our study, DCA levels were elevated in both feces and cecal homogenates from HFVHC- and HF-fed SPF mice (**Figure 4H, 5D**). We demonstrated that DCA activates LX-2 HSCs *in vitro*, with the most robust activation observed at physiologically relevant concentrations (10 µM) (**Figure 5E**). These findings suggest that DCA present in the gut may contribute to HSC activation following BA resorption back into circulation. Importantly, DCA is increased in patients with MASH^80–83^ and has been implicated in MASLD and MASH through mechanisms that include induction of pyroptosis and inflammation in hepatocytes^56,57^. Although our results support a role for DCA in activation of HSCs (**Figure 5B**) future studies are needed to test whether DCA alone is sufficient to promote hepatic fibrogenesis.

Beyond BA sensitivity, microbial shifts in response to HFVHC feeding observed in our study may also be explained in part by their ability to either assimilate, metabolize, or tolerate excess luminal cholesterol^26,27^. For example, *Eubacterium coprostanoligenes*, which encodes the intestinal sterol metabolism A (*ismA*) enzyme responsible for converting cholesterol to coprostanol in the gut^25^ exhibited cholesterol-dependent enrichment in mice fed LFVHC and HFVHC diet (**Figure S3A**). On the other hand, *B. pseudolongum* has been previously shown to assimilate dietary cholesterol to a high degree *in vivo*^26^. The *Bifidobacteria* ASV observed in our study that was depleted in response to cholesterol and saturated fat (**Figure S4**) mapped with ∼100% identity to *B. pseudolongum* (data not shown). Similarly, both glycosylation and dehydrogenation of cholesterol have been reported in several *Oscillibacter* members, a genus that also largely decreased in abundance in response to dietary cholesterol in our study (**Figure S3A, S4A**). Additional mechanistic studies, including those using gnotobiotic mice, will be required to determine how these interactions shape gut microbiota community membership and function and whether they contribute to the development of fibrosing MASH.

Our studies present some limitations that should be considered. First, all mice received glucose- and fructose-supplemented drinking water, limiting our capacity to assess specific interactions between other dietary components and gut microbes in the context of MASLD/MASH development and progression. Second, only male mice were used, which limits the generalizability of our findings regarding host-microbe-diet interactions to female mice, and hence, to human patient populations. Importantly, MASLD exhibits sex-dimorphic patterns, with lower risk in premenopausal women compared to men, but similar prevalence post-menopause due to complex gender-specific factors^84^. Third, we did not include a cohort of GF mice fed diet for 8 weeks, preventing direct comparisons between GF and SPF mice during early stages of disease. While our *in vitro* studies demonstrate that gut-derived factors from HFVHC-fed SPF mice are sufficient to activate HSCs, and we identified DCA as a potential mediator (**Figure 5, S5**), it is likely other BAs and microbial metabolites present in cecal homogenates also contribute to HSC activation. Lastly, although we provide an extensive characterization of diet-induced shifts in gut microbiota, the mechanisms driving these microbial shifts remain speculative and require further investigation.

In conclusion, our work highlights the synergistic effects of dietary cholesterol and saturated fat in shaping gut microbiota composition and their subsequent influence on hepatic fibrosis via gut-derived factors. Using a multifactorial design, we identified key aspects of host metabolic health, gut microbiota community membership, and fecal BA profiles that are differentially influenced by dietary components and microbial presence across early and late stages of MASLD/MASH. Our findings suggest that specific gut-derived factors, including DCA, other BAs, and microbial taxa, such as *Parasutterella*, may play a critical role in activating HSCs and promoting fibrogenesis in MASH. These insights provide a foundation for future studies aimed at elucidating microbial and metabolic pathways involved in hepatic fibrosis, with the goal of developing microbiota- and diet-based strategies to prevent or reverse MASH disease course.

## ACKNOWLEDGEMENTS

The authors would like to acknowledge the University of Wisconsin-Madison Gnotobiotic Facility staff, particularly Eugenio Vivas and Annelise Resende, as well as the Biochemistry/Biochemical Sciences Animal Facility staff, particularly Dustin Irving and Heather Morehouse, for animal husbandry support. We acknowledge the Dairy Innovation Hub Histology Resource and we are grateful to Laurie A. Sand for administrative support. We extend our thanks to Nancy Liu, Annabell Noel, Reagan Rodenbostel, Evan Silver, and Ella Blackburn for their technical support with *in vitro* and *in vivo* experiments.

## DATA AVAILABILITY STATEMENT

The data that support the findings of this study are available in the National Center for Biotechnology Information (NCBI) Sequence Read Archive (SRA) BioProject ID PRJNA1289062.

---

**Figure S1.**
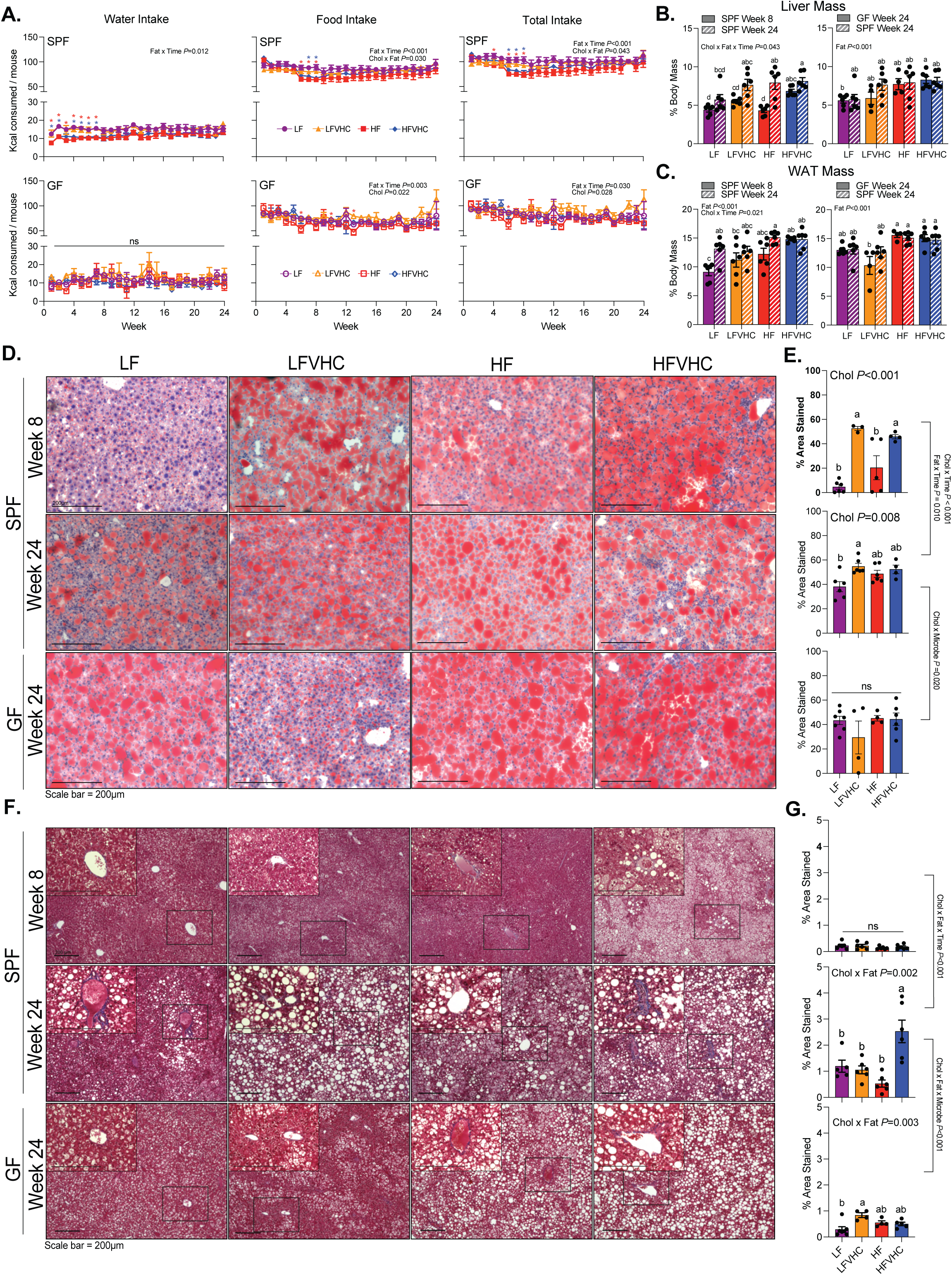
GF and SPF mice exhibit comparable caloric intake and adiposity, but disparate liver lipid and fibrosis profiles. **A)** Caloric intake attributed to water, food, or combined during the duration of the study. Data represent means ± SEM, analyzed via 4-way RM ANOVA (factors: Cholesterol, Fat, Microbes, Time) followed by 3-way ANOVA (factors: Cholesterol, Fat, Time) within SPF and GF groups and Tukey’s multiple comparisons within timepoint. \**P*<0.05. Asterisk color represents which group is significantly different from LF. **B,C)** Liver mass (B) and total white adipose tissue (WAT) mass (C) as a percent of body mass. Data represent means ± SEM analyzed via 3-way ANOVA. Comparisons made between SPF Week 8 vs. SPF Week 24 (factors: Cholesterol, Fat, Time, left panels) or between SPF Week 24 vs. GF Week 24 (factors: Cholesterol, Fat, Microbes, right panels), followed by Tukey’s multiple comparisons within timepoint. Bars with the same letter are not significantly different (*P*>0.05). **D)** Representative Oil Red O-stained liver sections; scale bar=200µm. **E)** Quantification of percent area stained red (indicating neutral lipids). **F)** Representative Masson’s Trichrome-stained liver sections; scale bar=200µm. Inset images (400x) correspond to boxed area in 100x images. **G)** Quantification of percent area stained blue (indicating collagen). Data represent means ± SEM, analyzed via 3-way ANOVA. Comparisons made between SPF Week 8 and SPF Week 24 (factors: Cholesterol, Fat, Time) or between SPF Week 24 and GF Week 24 (factors: Cholesterol, Fat, Microbes), followed by Tukey’s multiple comparisons. Bars with the same letter are not significantly different (*P*>0.05).

**Figure S2.**
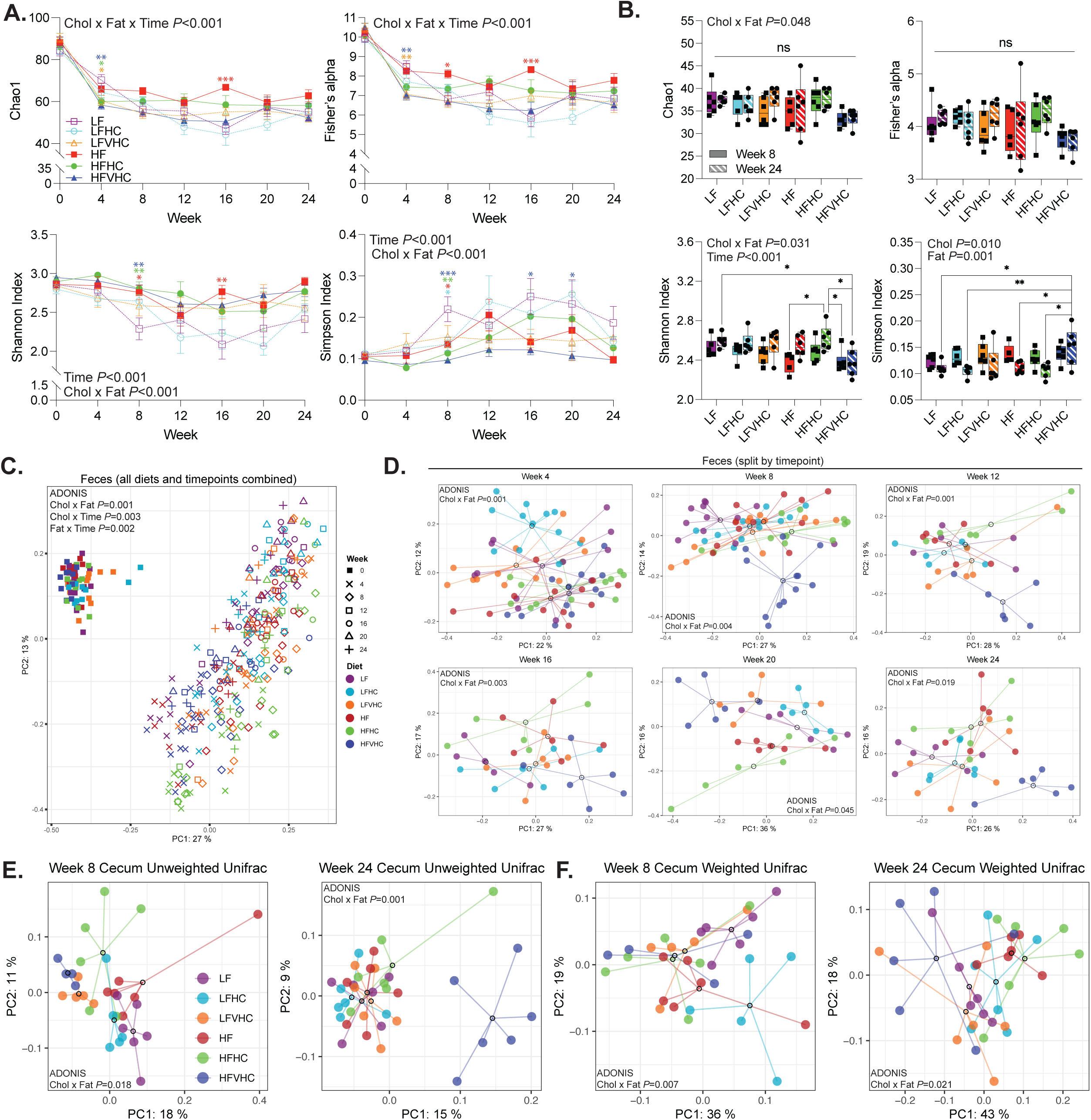
Dietary cholesterol and saturated fat differentially impact fecal and cecal microbiota diversity and composition over time in SPF mice. **A,B)** Chao1 (top left), Fisher’s alpha (top right), Shannon index (bottom left), and Simpson index (bottom right) α-diversity indices of fecal (A) and cecal (B) microbiota in SPF mice throughout the study (A). Data represent means ± SEM, analyzed via 3-way ANOVA (Factors: Cholesterol, Fat, Time) followed by Tukey’s multiple comparisons within timepoint. \**P*<0.05, \*\**P*<0.01, \*\*\**P*<0.005. For fecal diversity indices, asterisk color represents significant differences from LF. **C,D)** Bray-Curtis β-diversity PCoA of fecal microbiota, analyzed via 3-factor ADONIS (Factors: Time, Cholesterol, Fat) (C) or 2-factor ADONIS (Factors: Cholesterol, Fat) (D). **E,F)** Unweighted (E) and Weighted (F) Unifrac β-diversity PCoA of cecal microbiota after 8 (left panels) and 24 (right panels) weeks on diet, analyzed via 2-factor ADONIS (Factors: Cholesterol, Fat). For all PCoA plots, dots represent individual mice, open circles represent centroids with lines connecting individual dots within a treatment group.

**Figure S3.**
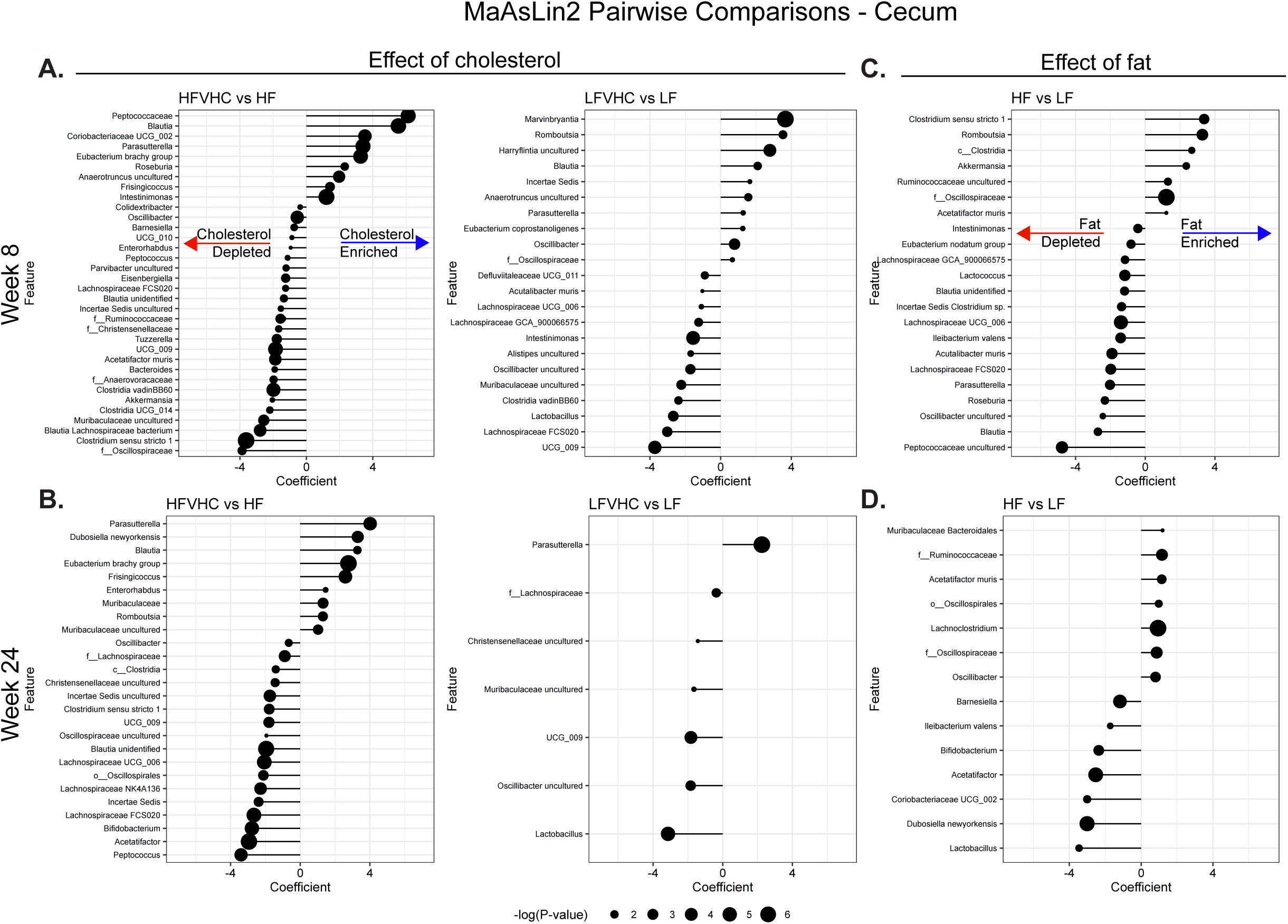
Distinct effects of dietary cholesterol and saturated fat on cecal microbiota composition at 8 and 24 weeks. **A-D)** ASVs significantly enriched or depleted by dietary cholesterol (A,B) or saturated fat (C,D) via MaAsLin2 pairwise comparisons at 8 (A,C) and 24 (B,D) weeks on diet. Positive coefficients indicate enrichment; negative coefficients indicate depletion. Dot size reflects -log(P-value). Taxonomic lables: “f__”: family-level annotation; “o__” order-level annotation; “c__” class-level.

**Figure S4.**
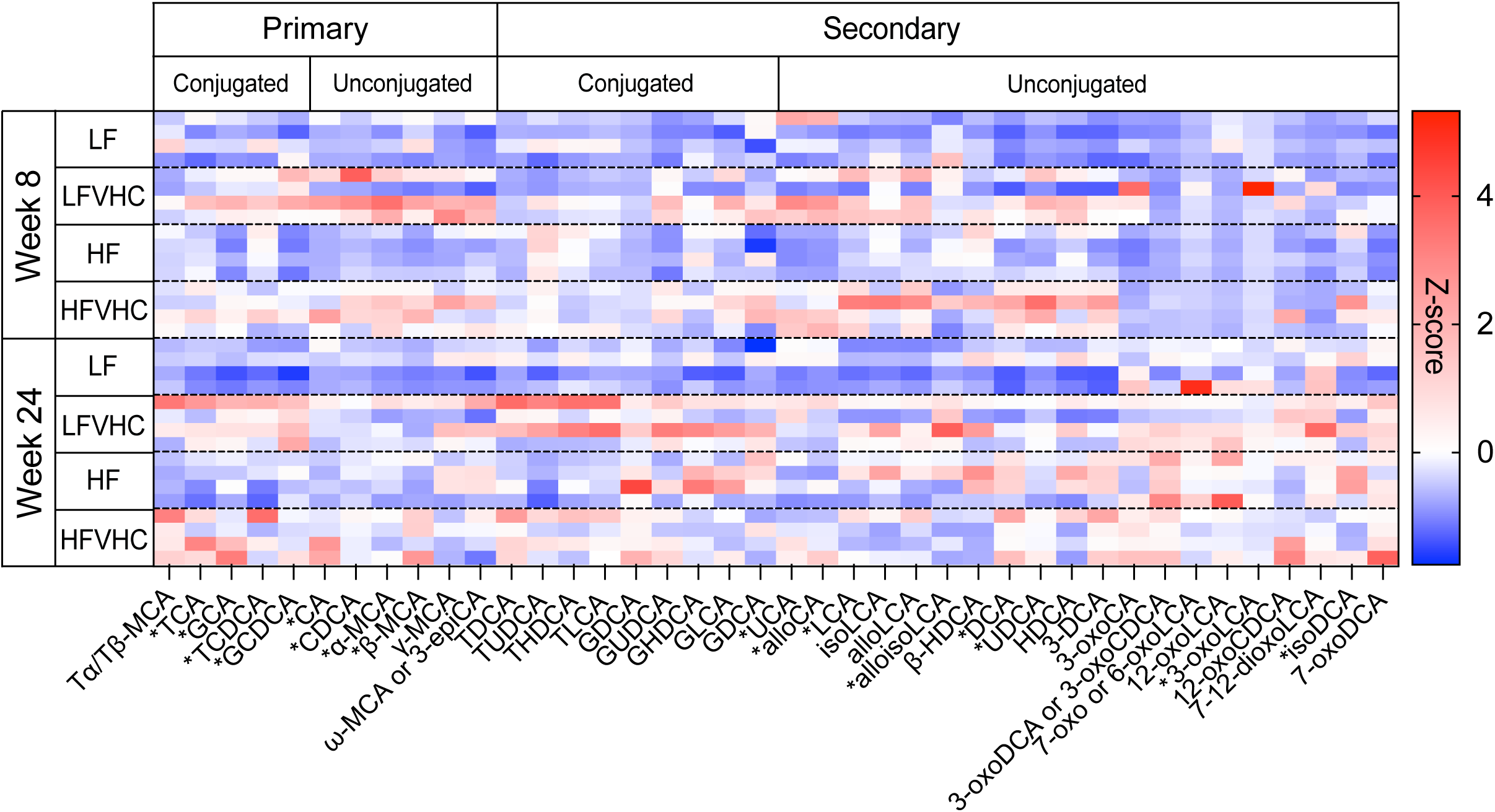
Fecal BA profiles are altered in response to dietary cholesterol and saturated fat. Heat map of fecal BA after 8 or 24 weeks in SPF mice, represented as Z-score of normalized peak area within each BA. BA with an asterisk(*) yielded quantitative values as shown in Figure 4G-L. Tα/β-MCA: Tauroα/β-muricholic acid; TCA: Taurocholic acid; GCA: Glycocholic acid; TCDCA: Taurochenodeoxycholic acid; GCDCA: Glycochenodeoxycholic acid; CA: Cholic acid; CDCA: Chenodeoxycholic acid; α/β/γ/ω-MCA: α/β/γ/ω-Muricholic acid; 3-epiCA: 3-epicholic acid; TDCA: Taurodeoxycholic acid; TUDCA: Tauroursodeoxycholic acid; THDCA: Taurohyodeoxycholic acid; TLCA: Taurolithocholic acid; GDCA: Glycodeoxycholic acid; GUDCA: Glycoursodeoxycholic acid; GLCA: Glycolithocholic acid; GDCA: Glycodeoxycholic acid; UCA: Ursocholic acid; alloCA: Allocholic acid; LCA: Lithocholic acid; isoLCA: Isolithocholic acid; alloLCA: Allolithocholic acid; alloisoLCA: Alloisolithocholic acid; β-HDCA: β-hyodeoxycholic acid; DCA: Deoxycholic acid; UDCA: Ursodeoxycholic acid; HDCA: Hyodeoxycholic acid; 3-DCA: 3-deoxycholic acid; 3-oxoCA: 3-oxocholic acid; 3-oxo(C)DCA: 3-oxochenodeoxycholic acid or 3-oxodeoxycholic; 7-/6-oxoLCA: 7-/6-oxolithocholic acid; 12-oxoLCA: 12-oxolithocholic acid; 3-oxoLCA: 3-oxolithocholic acid; 12-oxoCDCA: 12-oxochenodeoxycholic acid; 7-12-dioxoLCA: 7-12-dioxolithocholic acid; isoDCA: isodeoxyholic acid; 7-oxoDCA: 7-oxodeoxycholic acid.

**Figure S5:**
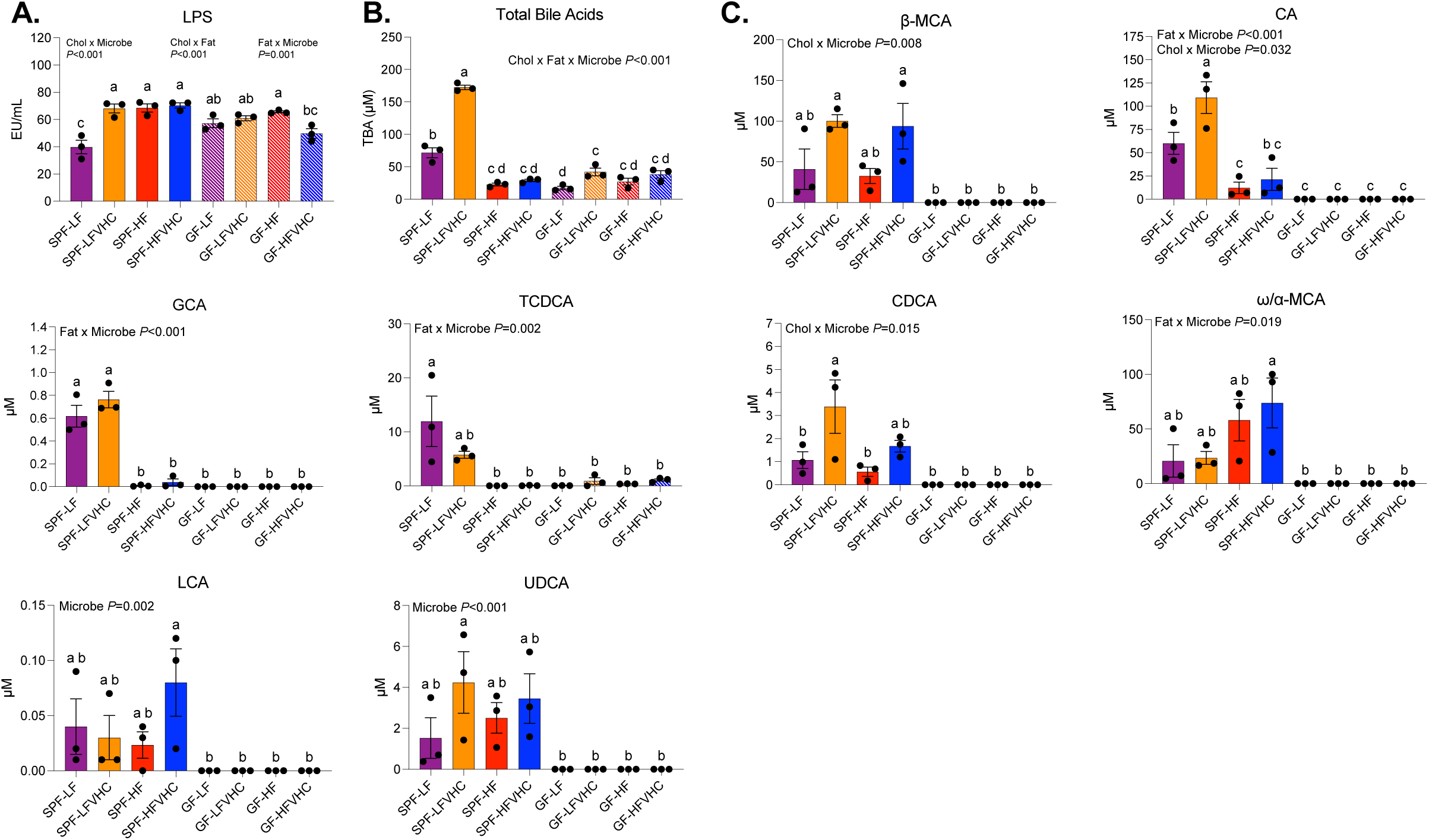
LPS and TBA are not primary mediators of *in vitro* HSC activation induced by cecal homogenates. **A,B)** LPS (A) and TBA concentrations (B) in cecal homogenates prepared as shown in Figure 5A. **C-G)** Individual BA concentrations in cecal homogenates. Data represent means ± SEM, analyzed via 3-way ANOVA (factors: Cholesterol, Fat, Microbes) followed by Tukey’s multiple comparisons within timepoint. Bars with the same letter are not significantly different (*P*>0.05). α/β/ω-MCA: α/β/ω-Muricholic acid; CA: Cholic acid; GCA: Glycocholic acid; TCDCA: Taurochenodeoxycholic acid; CDCA: Chenodeoxycholic acid; LCA: Lithocholic acid; UDCA: Ursodeoxycholic acid.

