## Supplemental Table 1 for "Gut microbes mediate the synergistic effects of dietary cholesterol and saturated fat in driving fibrosing MASH"

Table S1. Composition of semi-purified diets.

| Diet | Low-fat (LF) | Low-fat high-cholesterol (LFHC) | Low-fat very high-cholesterol (LFVHC) | High-fat (HF) | High-fat high-cholesterol (HFHC) | High-fat very high-cholesterol (HFVHC) |
| --- | --- | --- | --- | --- | --- | --- |
| Total energy (kcal/g) | 3.6 | 3.6 | 3.6 | 4.6 | 4.6 | 4.5 |
| Protein (% wt) | 17.3 | 17.3 | 17.3 | 17.3 | 17.3 | 17.3 |
| Carbohydrate (% wt) | 61.3 | 61.1 | 61.3 | 49.1 | 49.0 | 47.3 |
| Fat (% wt) | 5.2 | 5.2 | 5.2 | 21.2 | 21.2 | 21.1 |
| Milk Fat (% wt) | 4.0 | 4.0 | 4.0 | 21.0 | 21.0 | 21.0 |
| Cholesterol (% wt) | 0.05 | 0.20 | 2.00 | 0.05 | 0.20 | 2.00 |
| Glucose (18.1 g/L) | *✓* | | | | | |
| Fructose (23.1 g/L) | *✓* | | | | | |
