## Supplemental Table 2 for "Gut microbes mediate the synergistic effects of dietary cholesterol and saturated fat in driving fibrosing MASH"

Table S2. Primers used in this study.

| **Primer** | **Sequence (5’ 🡪 3’)** |
| --- | --- |
| **Mouse Primers** | |
| *Gapdh* (fwd) | CAAGGACACTGAGCAAGAGA |
| *Gapdh* (rvs) | TTGATGGTATTCAAGAGAGTAGGG |
| *Timp1* (fwd) | GGCATCCTCTTGTTGCTATC |
| *Timp1* (rvs) | CTTATGACCAGGTCCGAGTT |
| *Timp2* (fwd) | TAATTGCAGGAAAGGCAGAA |
| *Timp2* (rvs) | CTTCTTCTGGGTGATGCTAAG |
| *Mmp2* (fwd) | CTTCTTCAAGGACCGGTTTAT |
| *Mmp2* (rvs) | CCTCATACACAGCGTCAATC |
| *Mmp9* (fwd) | CACTGGGCTTAGATCATTCC |
| *Mmp9* (rvs) | AGCCACGACCATACAGATA |
| *Tgf* **β***1* (fwd) | CTATTGCTTCAGCTCCACAG |
| *Tgf* **β***1* (rvs) | GACAGAAGTTGGCATGGTAG |
| *Acta2* (fwd) | TCCCAGACATCAGGGAGTAA |
| *Acta2* (rvs) | CTATCGGATACTTCAGCGTCAG |
| *Col1a1* (fwd) | AGTCAGCAGATTGAGAACATCC |
| *Col1a1* (rvs) | ATCCAGTACTCTCCGCTCTT |
| *Igf1* (fwd) | TACTTCAACAAGCCCACAG |
| *Igf1* (rvs) | CATCTCCAGTCTCCTCAGAT |
| *Tlr4* (fwd) | AAATGCACTGAGCTTTAGTGGT |
| *Tlr4* (rvs) | TGGCACTCATAATGATGGCAC |
| *Il*-*1β* (fwd) | GAAATGCCACCTTTTGACAGTG |
| *Il*-*1β* (rvs) | TGGATGCTCTCATCAGGACAG |
| *Il-6* (fwd) | TCCTACCCCAATTTCCAATG |
| *Il*-*6* (rvs) | TTGCCGAGTAGATCTCAAAG |
| *Tnfα* (fwd) | CCTGTAGCCCACGTCGTAG |
| *Tnfα* (rvs) | GGGAGTAGACAAGGTACAACCC |
| *Cyp7a1* (fwd) | AAACTCCCTGTCATACCACAAAG |
| *Cyp7a1* (rvs) | TTTCCATCACTTGGGTCTATGC |
| *Cyp27a1* (fwd) | GAGGGCAAGTACCCAATAAG |
| *Cyp27a1* (rvs) | CTCTCCTTGTGCGATGAAG |
| **Human (LX-2) Primers** | |
| *GAPDH* (fwd) | GGAGCGAGATCCCTCCAAAAT |
| *GAPDH* (rvs) | GGCTGTTGTCATACTTCTCATGG |
| *COL1A1* (fwd) | AAAGGTGCTGATGGCTCTC |
| *COL1A1* (rvs) | GGACCACTTTCACCCTTGT |
| *IL-1β* (fwd) | TTCGACACATGGGATAACGAGG |
| *ACTA2* (fwd) | TCCTTCGTTACTACTGCTGA |
| *ACTA2* (rvs) | TCTTCTCAAGGGAGGATGAG |
| *IL-1β* (rvs) | TTTTTGCTGTGAGTCCCGGAG |
| *IL-6* (fwd) | CCTGAACCTTCCAAAGATGGC |
| *IL-6* (rvs) | TTCACCAGGCAAGTCTCCTCA |
| *MCP1* (fwd) | CAGCCAGATGCAATCAATGCC |
| *MCP1*(rvs) | TGGAATCCTGAACCCACTTCT |
| *TGFβR2* (fwd) | CACCTGTGACAACCAGAAA |
| *TGFβR2* (rvs) | CTCGTCATTCTTTCTCCATACA |
| *TNFα* (fwd) | GAGGCCAAGCCCTGGTATG |
| *TNFα* (rvs) | CGGGCCGATTGATCTCAGC |
| **Bacterial 16S rRNA Gene Amplicon Primers** | |
| 515F (Caporaso) | GTGCCAGCMGCCGCGGTAA |
| 806R (Caporaso) | GGACTACHVGGGTWTCTAAT |
