## Supplemental Table 3 for "Gut microbes mediate the synergistic effects of dietary cholesterol and saturated fat in driving fibrosing MASH"

Table S3. Bile acid names and structure description.

| **Bile Acid Common Name** | **Abbreviation** | **Mol. Formula** | **C1** | **C2** | **C3** | **C6** | **C7** | **C12** | **R** | **Rt** |
| --- | --- | --- | --- | --- | --- | --- | --- | --- | --- | --- |
| Glycocholic acid | GCA | C26H43NO6 | α,β-H | α,β-H | α-OH | α,β-H | α-OH | α-OH | G | 14.05 |
| Cholic acid | CA | C24H40O5 | α,β-H | α,β-H | α-OH | α,β-H | α-OH | α-OH | H | 15.04 |
| Taurochenodeoxycholic acid | TCDCA | C26H45NO6S | α,β-H | α,β-H | α-OH | α,β-H | α-OH | α,β-H | T | 15.73 |
| Chenodeoxycholic acid | CDCA | C24H40O4 | α,β-H | α,β-H | α-OH | α,β-H | α-OH | α,β-H | H | 17.1 |
| Ursodeoxycholic acid | UDCA | C24H40O4 | α,β-H | α,β-H | α-OH | α,β-H | β-OH | α,β-H | H | 13.75 |
| Lithocholic acid | LCA | C24H40O3 | α,β-H | α,β-H | α-OH | α,β-H | α,β-H | α,β-H | H | 19.09 |
| α-muricholic acid | α-MCA | C24H40O5 | α,β-H | α,β-H | α-OH | β-OH | α-OH | α,β-H | H | 11.83 |
| β-muricholic acid | β-MCA | C24H40O5 | α,β-H | α,β-H | α-OH | β-OH | β-OH | α,β-H | H | 12.03 |
| ω-muricholic acid | ω-MCA | C24H40O5 | α,β-H | α,β-H | α-OH | α-OH | β-OH | α,β-H | H | 11.89 |
